# Scaffold subunits support associated subunit assembly in the *Chlamydomonas* ciliary nexin-dynein regulatory complex

**DOI:** 10.1101/684316

**Authors:** Long Gui, Kangkang Song, Douglas Tristchler, Raqual Bower, Yan Si, Aguang Dai, Katherine Augsperger, Jason Sakizadeh, Magdalena Grzemska, Thomas Ni, Mary E. Porter, Daniela Nicastro

## Abstract

The nexin-dynein regulatory complex (N-DRC) in motile cilia and flagella functions as a linker between neighboring doublet microtubules, acts to stabilize the axonemal core structure, and serves as a central hub for the regulation of ciliary motility. Although the N-DRC has been studied extensively using genetic, biochemical, and structural approaches, the precise arrangement of the eleven (or more) N-DRC subunits remains unknown. Here, using cryo-electron tomography, we have compared the structure of *Chlamydomonas* wild-type flagella to that of strains with specific DRC subunit deletions or rescued strains with tagged DRC subunits. Our results show that DRC7 is a central linker subunit that helps connect the N-DRC to the outer dynein arms. DRC11 is required for the assembly of DRC8, and DRC8/11 form a sub-complex in the proximal lobe of the linker domain that is required to form stable contacts to the neighboring B-tubule. Gold labeling of tagged subunits determines the precise locations of the previously ambiguous N-terminus of DRC4 which is now shown to contribute to the core scaffold of the N-DRC and C-terminus of DRC5. Our results reveal the overall architecture of N-DRC, with the three subunits, DRC1/2/4 forming a core complex that serves as the scaffold for the assembly of the “functional subunits” associate, namely DRC3/5-8/11. These findings shed light on N-DRC assembly and its role in regulating flagellar beating.

**Significance Statement:** Cilia and flagella are small hair-like appendages in eukaryotic cells that play essential roles in cell sensing, signaling, and motility. The highly conserved nexin-dynein regulatory complex (N-DRC) is one of the key regulators for ciliary motility. At least 11 proteins (DRC1–11) have been assigned to the N-DRC, but their precise arrangement within the large N-DRC structure is not yet known. Here, using cryo-electron tomography combined with genetic approaches, we have localized DRC7, the sub-complex DRC8/DRC11, the N-terminus of DRC4, and the C-terminus of DRC5. Our results provide insights into the N-DRC structure, its function in the regulation of dynein activity, and the mechanism by which *n-drc* mutations can lead to defects in ciliary motility that cause disease.

## INTRODUCTION

Cilia and flagella are dynamic microtubule (MT)-based organelles that emanate from the surface of many eukaryotic cells and are involved in cell sensory functions, motility, and signaling. Defects in cilia assembly or function have been associated with multiple human disorders collectively known as ciliopathies, such as polycystic kidney disease, Bardet-Biedl syndrome, infertility, hydrocephalus, and primary ciliary dyskinesia (1, 2).

The microtubule-based axoneme forms the core structure of motile cilia and is highly conserved, from the green algae *Chlamydomonas reinhardtii* to differentiated cells in the human body. The “9 + 2” axoneme is comprised of nine outer doublet microtubules (DMTs) and a central-pair complex (CPC) composed of two singlet microtubules and associated projections (Fig. 1*A*). Each DMT is built from many copies of a 96 nm long unit that repeats along the length of the axoneme and consists of an A-tubule and B-tubule. The outer and inner dynein arms (ODAs and IDAs, respectively) are arranged in two distinct rows along the length of the A-tubule of each DMT and function as the motors that drive ciliary motility. The dynein motors “walk” on the B-tubule of the neighboring DMT and thereby generate sliding forces between the doublets (3). The nexin link between DMTs is thought to restrict this inter-doublet sliding and thus transform sliding into axonemal bending (4–6). Our previous ultrastructural study demonstrated that the nexin link is an integral part of the dynein regulatory complex, referred to as the nexin-dynein regulatory complex (N-DRC) (7). Formation of ciliary bending in different directions and beating patterns require precise regulation and coordination of the activities of thousands of axonemal dyneins (8–10). Genetic, biochemical and structural studies suggest that several complexes are involved in the transduction of signals that ultimately control the activity of the dyneins. These regulatory complexes include the CPC, the radial spokes, the calmodulin- and radial spoke-associated complex (CSC), the I1 inner dynein (or dynein f) on the proximal end of the 96-nm axonemal repeat, and the N-DRC on the distal end of the repeat (11).

**Fig. 1.**
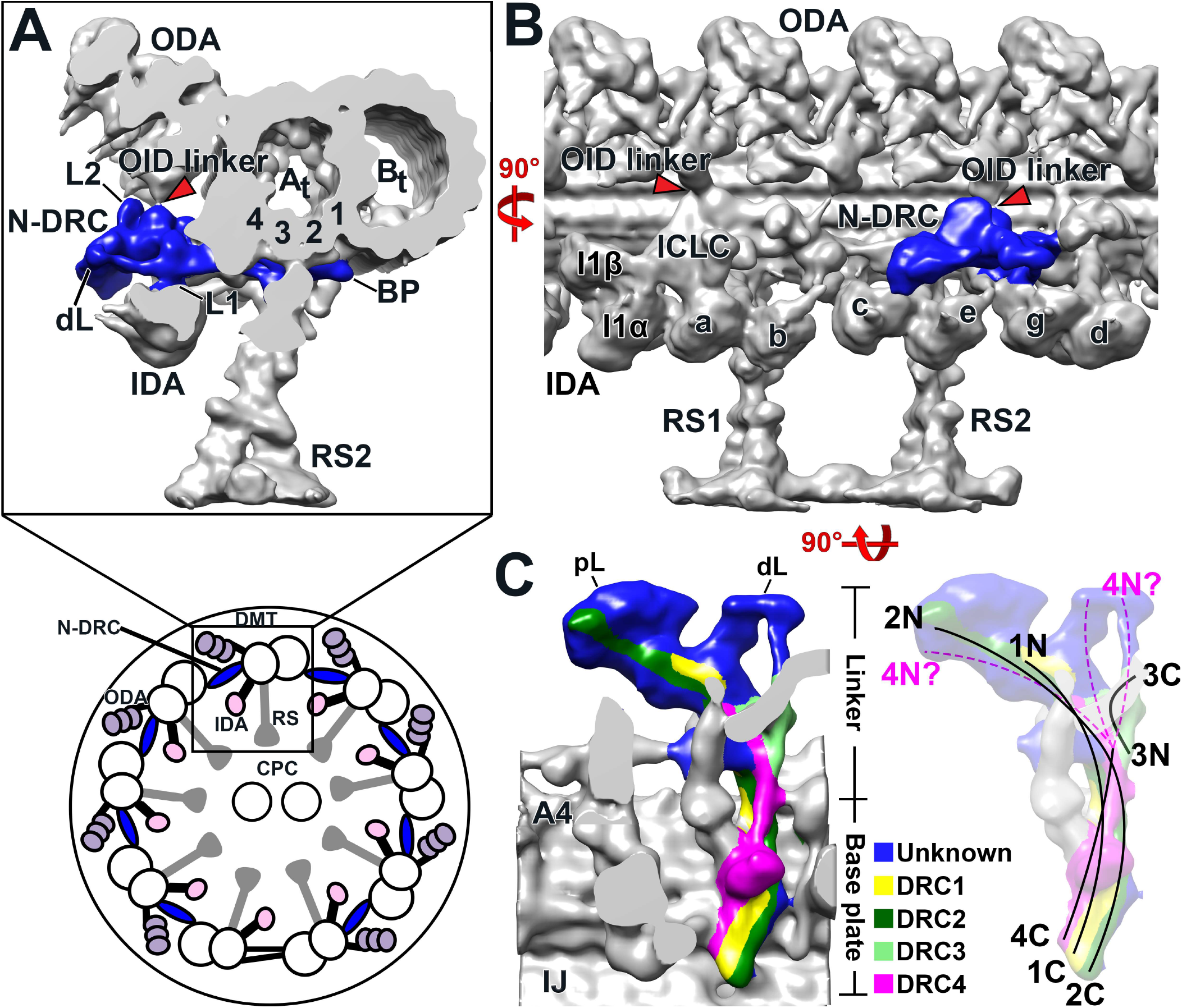
Structure of the N-DRC and organization of known N-DRC subunits in *Chlamydomonas* flagella. **(A, bottom)** Schematic of a *Chlamydomonas* flagellum in cross-sectional view, seen from the flagellar tip toward the base. The outer and inner dynein arms (ODA, IDA) and the nexin-dynein regulatory complex (N-DRC) connect the neighboring doublet microtubules (DMTs), whereas the radial spokes (RS) connect to the central pair complex (CPC). **(A top, B, and C)** Isosurface rendering of the three-dimensional structure of the 96-nm axonemal repeat in *Chlamydomonas* wild-type flagella reconstructed by cryo-electron tomography (cryo-ET), shown in cross-sectional (*A top*), longitudinal front (*B*) and bottom views (*C*). (*A top and B*) overviews showing the location of the N-DRC (blue) within the axonemal repeat, and connections between the N-DRC and other axonemal structures, including the outer dynein-inner dynein (OID) linkers (red arrowheads). The N-DRC linker regions has 3 protrusions: the L1- and L2 protrusions, and the OID linker. (*C*) 3D maps of the N-DRC in longitudinal views from the bottom. The known locations of DRC1-4 (1–4) are colored; N and C correspond to the N- and C-termini of the proteins, respectively. The dashed line indicates ambiguity in localization of the N-terminal region of DRC4. Note that DRC5 and DRC6 are thought to be located at the L2 projection, which is in the dorsal side of the linker and thus cannot be visualized in (*C*). Other labels: At, A-tubule (protofilaments 1-4 are labelled in panel A top, and A4 in panel C); B_t_, B-tubule; BP, base plate; dL, distal lobe; pL, proximal lobe; IJ, inner junction; *a–e* and *g*, inner dynein arm isoforms; I1α/ I1β/ ICLC, α- and β-heavy chain, and intermediate-light chain complex of I1 dynein.

The N-DRC is a large 1.5-MDa structure consisting of two main domains, the base plate and the linker (Fig. 1*C*). The base plate attaches to the DMT, beginning at the inner junction between the A- and B-tubule and ending at protofilament A4. The linker domain extends across the inter doublet space to the neighboring B-tubule (7). Genetic and biochemical studies have demonstrated that the N-DRC is composed of at least 11 subunits, but the precise arrangement of these subunits in the NDRC structure remains elusive (12, 13). Recent work with tagged subunits has revealed the approximate locations of DRC1-6. The C-termini of DRC1, DRC2(FAP250), and DRC4 are located close to each other at the end of the base plate near the inner junction, and their N-termini extend through the linker toward the neighboring B-tubule. DRC3(FAP134), DRC5(FAP155) and DRC6(FAP169) are located in the linker region (7, 14–16). However, little is yet known about the locations of DRC7-11 or the specific interactions within the structure of the N-DRC. The identification of DRC7-11 mutants will be important for the dissection of the N-DRC and its function as regulatory hub.

Here, we have integrated genetic and biochemical approaches with cryo-electron tomography (cryo-ET) to localize DRC7 (FAP50) and the subcomplex DRC8/11 (FAP200/FAP82) within the N-DRC. DRC7 is found in the central region of the N-DRC linker, including the outer-dynein-inner-dynein (OID)-linker between the N-DRC and the row of outer dynein arms (Fig. 1*B*). The *drc11* mutant lacks two N-DRC subunits: DRC11 and DRC8, which localize to the proximal lobe of the N-DRC linker domain. We also use cryo-ET of SNAP-tagged DRCs to precisely locate the C-terminus of DRC5 in the middle region of the linker, and the N-terminus of DRC4 to the proximal lobe of the linker domain.

## RESULTS

### Identification of *drc* mutants for DRC7 and DRC11

The N-DRC contains at least eleven DRC subunits with distinct functional domains, but *drc* mutations have only been characterized in five genes (*drc1-drc5*) in *Chlamydomonas*. To extend the repertoire of *drc* mutants, we analyzed a collection of mutants generated by insertional mutagenesis the Chlamydomonas Library Project (CLiP) (17, 18) (see SI Materials and Methods). PCR confirmed the sites of plasmid insertion in two *drc7* strains and two *drc11* strains; however, the impact of plasmid insertion was highly variable (Table S1 and Fig. S1).

None of the *drc7* strains examined displayed an obvious motility defect by phase contrast light microscopy (Fig. 2*B* and Table S1), and so axonemes were isolated from two strains and analyzed on Western blots. No DRC7 band was detected in the 197909 strain (from now on referred to as *drc7*) (Fig. 2*C*), whereas strain 080526 showed two DRC7 related bands (Fig. S1*D*) that are likely the result of alternative splicing. To determine if the absence of DRC7 affected the assembly of other axonemal proteins, the *drc7* axonemes were labeled with isobaric tags for relative and absolute quantitation (iTRAQ) and analyzed by tandem mass spectrometry (MS/MS). Between about 500-650 proteins were identified by at least five peptides in two independent iTRAQ experiments. However, only one protein, DRC7, was significantly reduced (P value <0.05) below 50% in both experiments (Table 1). All the other DRC subunits were present at wild-type levels (Table 1), as confirmed by Western blots (Fig. 2*C*).

**Fig. 2.**
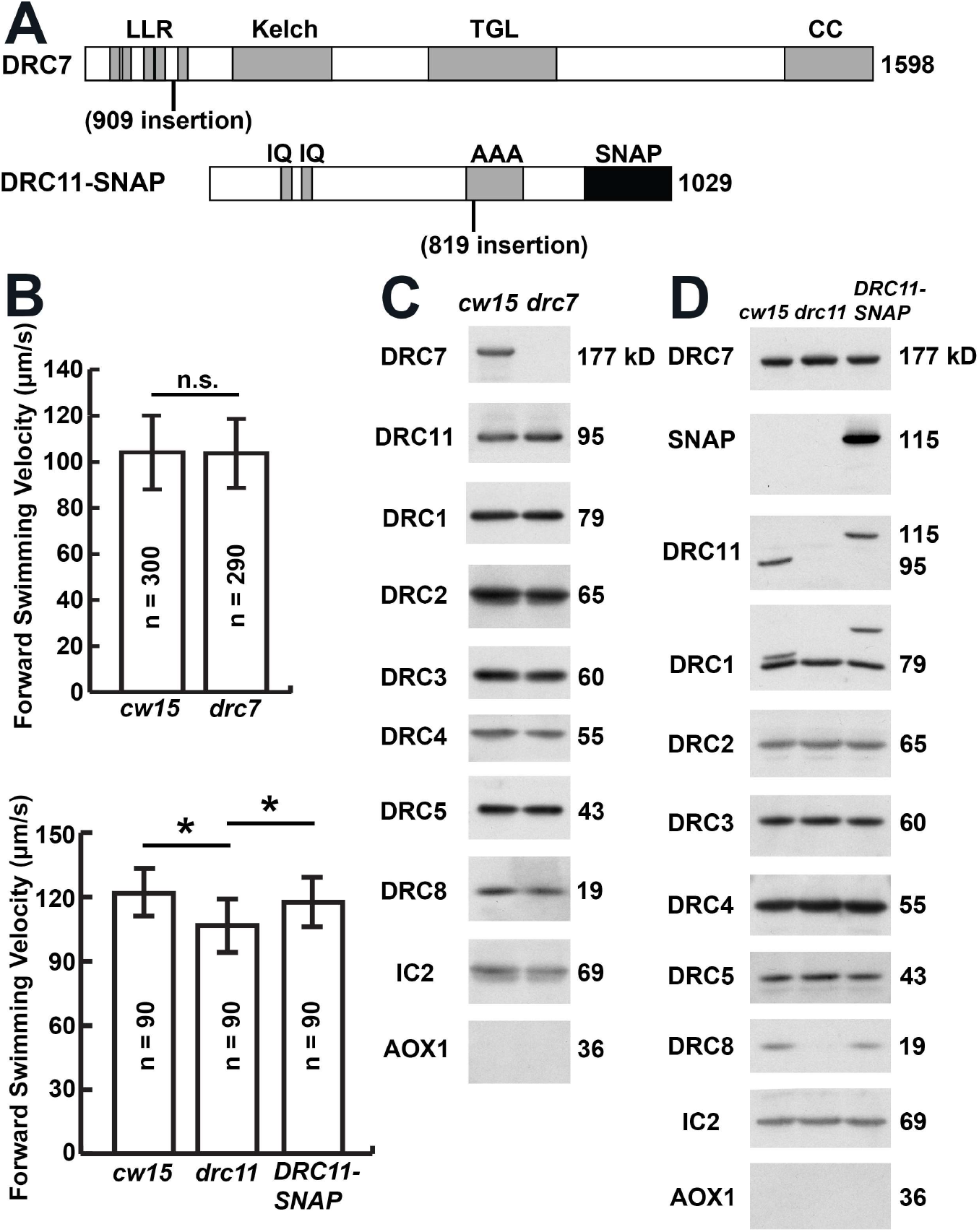
Identification of new mutations in *DRC7* and *DRC11*, and their impact on flagellar motility and assembly of the N-DRC. **(A)** The DRC7 and DRC11 protein sequences are drawn to scale, with the location of predicted polypeptide domains indicated: AAA, <u>A</u>TPase <u>a</u>ssociated with diverse cellular activities domain; CC, coiled-coil domain; IQ, calmodulin binding domain; Kelch, Kelch domain; LRR, leucine-rich repeat; SNAP, C-terminal SNAP tag; TGL, transglutaminase-like domain. Also shown are the approximate locations of the plasmid insertion sites in the *drc7* and *drc11* mutants. **(B)** The forward swimming velocities of *drc7* and *drc11* as measured by phase contrast microscopy are shown relative to the background strain (*cw15*). No significant difference was detected between *drc7* and *cw15*. The *drc11* strain was slightly slower (P < 0.05) than *cw15*, and transformation with a tagged WT gene, *DRC11-SNAP*, increased swimming velocities to near *cw15* levels. **(C, D)** Western blots of axonemes isolated from the “wild-type” background (*cw15*), the *drc* mutants, and the *DRC11-SNAP* rescued strain were probed with antibodies against several DRC subunits. Note the presence of a band detected by both the DRC11 and SNAP antibodies that migrated at the size predicted for a SNAP-tagged DRC11 subunit in (D). Antibodies against the IC2 subunit of the outer dynein arm served as a loading control, and antibodies against AOX1 served as a marker for cell body contamination.

**Table 1.** iTRAQ protein ratios in WT (cw-15) and *drc7* mutant axonemes.

| Protein | Experiment 1 |  |  | Experiment 2 |  |  |
| --- | --- | --- | --- | --- | --- | --- |
|  | Peptides | WT/WT | *drc7*/WT | Peptides | WT/WT | *Drc11*/WT |
| **DRC subunits** |  |  |  |  |  |  |
| DRC1 | 74 (137) | 1.09 | 1.17 | 77 (118) | 1.00 | 0.99 |
| DRC2 | 42 (169) | 1.10 | 1.25 | 39 (68) | 1.00 | 1.00 |
| DRC3 | 82 (81) | 1.06 | 1.10 | 72 (82) | 1.01 | 0.95 |
| DRC4 | 97 (106) | 1.06 | 1.16 | 86 (131) | 1.00 | 1.00 |
| DRC5 | 34 (24) | 1.04 | 0.95 | 39 (29) | 1.01 | 0.95 |
| DRC6 | 28 (29) | 1.04 | 0.88 | 27 (34) | 1.01 | 0.85 |
| **DRC7\*** | 103 (128) | 1.10 | **0.15** | 112 (152) | 1.02 | **0.15** |
| DRC8 | 14 (23) | 1.08 | 1.00 | 11 (25) | 0.98 | 0.95 |
| DRC9 | 41 (58) | 1.08 | 0.96 | 42 (56) | 1.01 | 0.89 |
| DRC10 | 37 (48) | 1.07 | 0.98 | 33 (51) | 1.02 | 0.91 |
| DRC11 | 90 (111) | 1.06 | 1.05 | 84 (121) | 1.00 | 1.00 |
| **MT doublet proteins** |  |  |  |  |  |  |
| Tektin | 109 (107) | 1.25 | 0.93 | 144 (135) | 0.99 | 1.11 |
| FAP59 | 83 (89) | 1.10 | 1.17 | 72 (96) | 1.00 | 0.99 |
| FAP172 | 95 (101) | 1.07 | 1.18 | 90 (112) | 1.02 | 1.07 |
| Rib43 | 80 (88) | 1.08 | 1.13 | 67 (84) | 1.00 | 1.01 |
| Rib72 | 241 (182) | 1.05 | 1.11 | 208 (202) | 0.98 | 0.97 |
| TUA | 701 (1018) | 1.03 | 1.07 | 588 (963) | 1.00 | 1.01 |
| TUB | 880 (1229) | 0.97 | 0.94 | 868 (1283) | 1.00 | 1.00 |

The amount of protein in each sample was compared to that present in WT (*cw-15*) to obtain a relative protein ratio. The WT/WT ratios indicate the variability in iTRAQ labeling and protein loading between replicates (typically less than 10%). For each iTRAQ experiment, duplicate mutant/WT ratios were averaged. Those proteins whose ratios were less than 50% of WT and statistically significant (P value < 0.05) in both experiments are shown in bold. Peptides indicate the number used for protein ID (95% confidence) followed by the number used for iTRAQ quantification (in parenthesis).

Although both *drc11* candidates had confirmed plasmid insertions in introns (Fig S1*E-G*), they displayed different motility phenotypes. The 130937 strain displayed an obvious slow swimming phenotype (Fig. S2*B*) and failed to assemble both DRC11 and DRC8 (Fig. S2*C*). However, transformation with a BAC clone (39g16) containing the *DRC11* gene failed to rescue the motility defect (0 rescues out of 538 transformants). This observation suggested the possibility of a second motility mutation in this strain. Further analysis by iTRAQ labeling, MS/MS, Western blots and cryo-ET revealed that the *drc11*-937 strain contained an unmapped mutation in a gene that disrupted the assembly of the outer dynein arms (Fig. S2*C-I* and Table S6).

The second *drc11* strain, 068819, displayed a slight but significant decrease in its forward swimming velocity (Fig. 2*B*). Western blots of axonemes confirmed that both DRC11 and DRC8 were missing, but several other DRC subunits and the outer dynein arms (as detected by an IC2/IC69 antibody) were present at wild-type levels (Fig. 2*D*). To determine if the loss of DRC8 and DRC11 affected the assembly of other polypeptides, the 068819 axonemes were analyzed by iTRAQ labeling and MS/MS. Between about 530-630 proteins were identified by at least five peptides in two independent experiments. However, only three proteins were significantly reduced below 30% in both iTRAQ experiments (Table 2): DRC8, DRC11, and Cre12.g513650. Little is known about Cre12.g513650; it is a small protein (146 amino acids) only found in green alga.

**Table 2.**
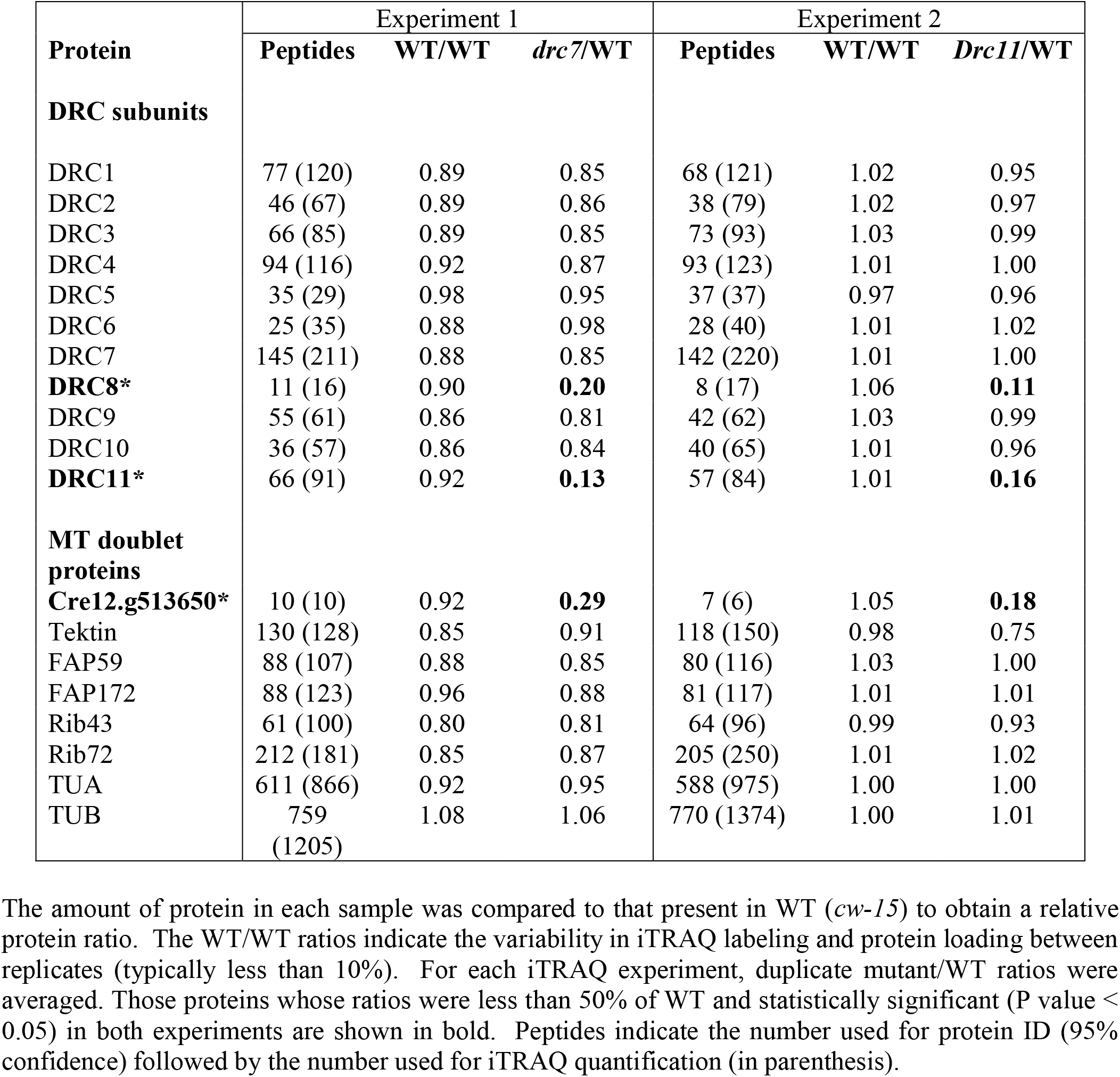
iTRAQ protein ratios in WT (cw-15) and *drc11* mutant axonemes.

To verify that the defects observed in the 068819 strain were caused by the *drc11* mutation, we transformed the mutant with a wild-type *DRC11* cDNA containing a C-terminal SNAP tag (Fig. 2*A*). A SNAP positive strain was recovered that increased forward swimming velocities to near wild-type levels (Fig. 2*B*) and restored the assembly of both DRC11 and DRC8, as shown by Western blots of axonemes (Fig. 2*D*). Furthermore, the DRC11-C-SNAP polypeptide migrated at the size expected for the SNAP tagged protein (~115 kDa). These results demonstrate that the 068819 strain is a bona fide *drc11* mutation and that DRC8 depends on the presence of DRC11 for its assembly into the axoneme.

### DRC7 localizes to the N-DRC linker including the OID-linker and distal lobe

Three-dimensional reconstruction of the axoneme by cryo-ET and sub-tomogram averaging revealed that the N-DRC linker begins approximately at protofilament A4 and projects into space between neighboring doublets (Fig. 1*A* and *C*) (7, 19). The linker domain contains several protrusions: L1 that connects to IDA g, L2 at the branch point between the proximal and distal lobes, and the OID-linker that connects the N-DRC to the ODAs (Fig. 1*A*). The proximal lobe is at least twice as the size of the distal lobe, but they both connect to the neighboring B-tubule (Fig. 1*C*).

To determine the location of DRC7 subunit, we compared the three-dimensional structure of wild-type and *drc7* axonemes by cryo-ET and sub-tomogram averaging (Fig. 3). We found missing densities only within the N-DRC linker domain and not in the base plate. Specifically, the *drc7* N-DRC lacks the OID-linker, a density on the distal side of the linker, and the most distal portion of the distal lobe (Fig. 3*D-F, J-L* and Movie S1), which reduces the size of the distal lobe interface with the neighboring B-tubule. The total mass of the densities missing in the *drc7* N-DRC is estimated to be ~200 kDa, suggesting that the density contains a single copy of DRC7, which has a molecular weight of 177 kDa.

**Fig. 3.**
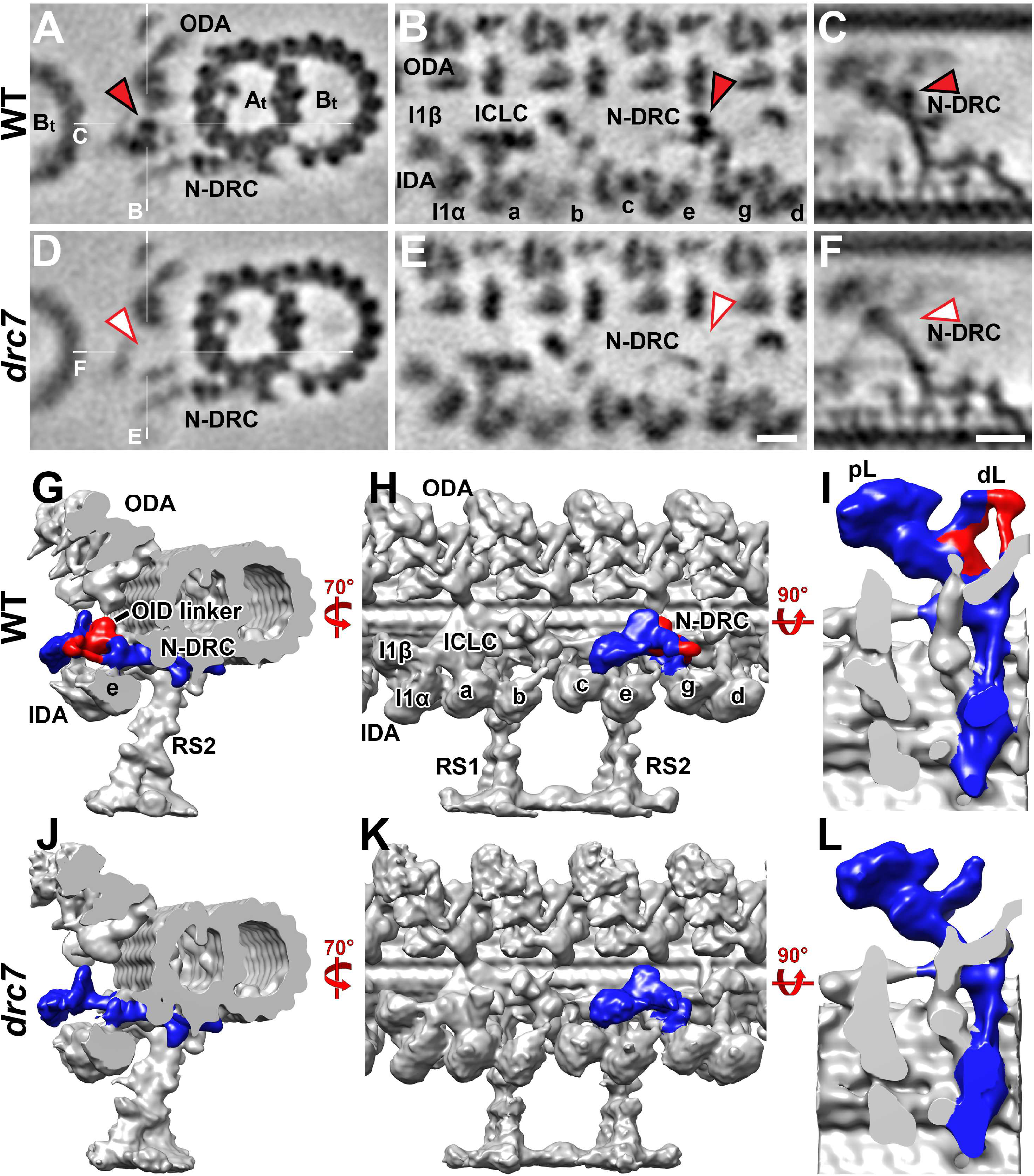
Comparison between wild-type and *drc7* axonemes reveals the location of DRC7 near the OID linker and in part of the distal lobe of the N-DRC. **(A–F)** Tomographic slices of the averaged 96-nm-long repeats from *Chlamydomonas* wild-type (A–C) and *drc7* axonemes (D–F) viewed in crosssectional (A and D) and longitudinal (B, C, E, and F) orientations. White lines indicate the locations of the slices in the respective panels. Electron densities corresponding to DRC7 in the nexin-dynein regulatory complex (N-DRC) of WT (red arrowheads in A-C) were absent from *drc7* axonemes (white arrowheads in D-F). **(G-L)** Isosurface renderings show the three-dimensional structures of the averaged axonemal repeats of wild-type (G-I) and *drc7* (J-L) in cross-sectional (G and J), longitudinal (H and K), and enlarged longitudinal bottom views (I and L; looking from the axoneme center outward). The structural difference between the WT and *drc7* is colored red in (G-I) and include the OID linker, a portion of the distal lobe (dL), and the connection between the distal lobe and the L1 protrusion (named connection #7 in (7)). Other labels: At, A-tubule; B_t_, B-tubule; IDA, inner dynein arm; ODA, outer dynein arm; RS, radial spoke; pL, proximal lobe; a–e and g, inner dynein arm isoforms; I1α/ I1β/ ICLC, α- and β-heavy chain, and intermediate-light chain complex of I1 dynein. Scale bar: 10 nm (in E, valid also for A, B, D), 10 nm (in F, valid also for C).

### DRC8/11 localize to the proximal lobe of the N-DRC linker

Our biochemical assays have indicated that the *drc11* strain lacks two polypeptides, DRC8 and DRC11. To determine the location of the DRC8/11 sub-complex, we analyzed *drc11* axonemes by cryo-ET and sub-tomogram averaging (Fig. 4). Compared with wild type, the averaged 96-nm repeats of *drc11* axonemes showed deficiencies in the proximal lobe of the N-DRC linker region (Fig. 4*D-F, J-L* and Movie S2). The structural defect greatly reduces the interface between the proximal lobe and the neighboring B-tubule. The missing density is estimated to be ~250 kDa. The morphology of the other N-DRC domains, including the distal lobe and the base plate, resembled the wild-type structure. Although we found that strain *drc11-937* was a double mutant with additional ODA defects, we were still able to detect the same N-DRC structural defects in this double mutant (Fig. S2*C-I*) as described for *drc11*.

**Fig. 4.**
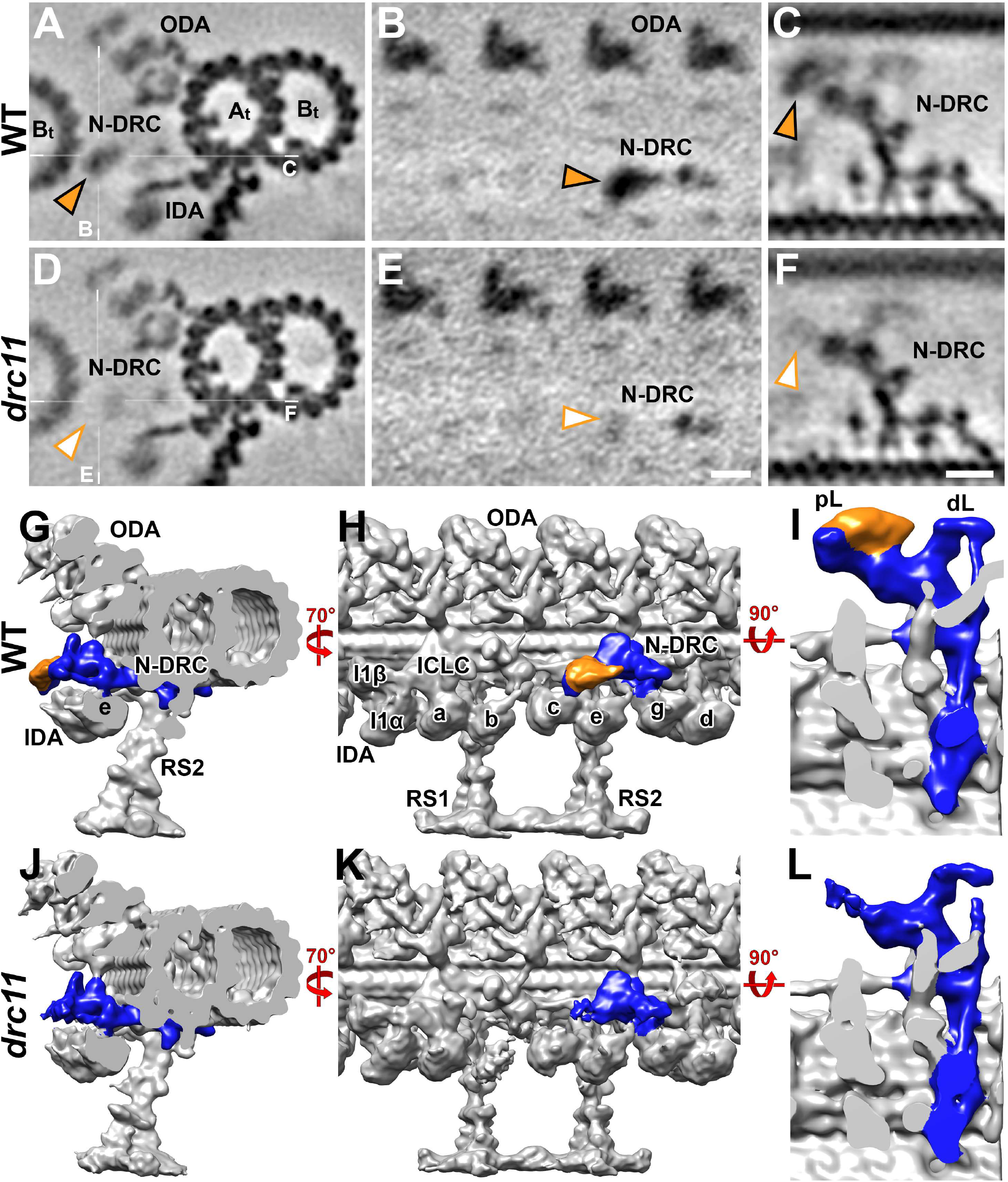
Three-dimensional localization of the DRC8/11 sub-complex at the proximal lobe of the N-DRC linker using cryo-ET. **(A–F)** Tomographic slices of the averaged 96-nm repeats of *Chlamydomonas* wild-type (A–C) and *drc11* axonemes (D–F) viewed in cross-sectional (A and D) and longitudinal (B, C, E, and F) orientations. White lines indicate the locations of the slices in the respective panels. Electron densities corresponding to DRC8/11 in the nexin-dynein regulatory complex (N-DRC) of WT (orange arrowheads in A-C) were missing from the *drc11* axonemes (white arrowheads in D-F). **(G-L)** Isosurface renderings show the 3D structures of the averaged axonemal repeats of wild-type (G-I) and *drc11* (J-L) in cross-sectional (G and J), longitudinal (H and K), and enlarged longitudinal bottom views (I and L; looking from the axoneme center outward). The structural difference between the WT and *drc11* is colored orange in (G-I) and includes a major portion of the proximal lobe (pL). Other labels: At, A-tubule; Bt, B-tubule; IDA, inner dynein arm; ODA, outer dynein arm; RS, radial spoke; dL, distal lobe; a–e and g, inner dynein arm isoforms; I1α/ I1β/ ICLC, α- and β-heavy chain, and intermediate-light chain complex of I1 dynein. Scale bar: 10 nm (in E, valid also for A, B, D), 10 nm (in F, valid also for C).

The tip region (close to the neighboring DMT) and the upper part of the proximal lobe (facing the ODA) were completely missing in *drc11* axonemes. In addition, we observed a remaining, yet weakened and blurred electron density that appears to be the lower portion of the proximal lobe that faces the IDAs (Fig. 5*B, G*). This blurring effect could be due to reduced occupancy or positional flexibility of the structures that comprise this region. Therefore, we applied automatic image classification analyses with different masks that focused the analyses on specific structures of interest (20). Classification of wild-type repeats showed one structurally homogenous state across all the axonemal repeats (Fig. 5*A* and *F*). In contrast, we found that the lower proximal lobe structure in *drc11* fell into four conformational classes, which differed mainly in its position (“height”) between the ODA and IDA rows (Fig. 5 and Movie S3). More specifically, 23% of the repeats showed the remaining proximal lobe density in a low position close to the IDAs (class 1, Fig. 5*C* and *H*); 25% were positioned in the middle (class 2, Fig. 5*D* and *I*); and 27% were located in a higher position closer to the ODAs (class 3, Fig. 5*E* and *J*). The remaining 25% could not be categorized because of blurred densities and anisotropic density distribution caused by the missing wedge-artifact typical for raw single-axis tomographic reconstructions (21). Thus, the proximal lobe of the N-DRC is composed of two parts: The DRC8/11 sub-complex comprises the tip and upper (ODA-facing) region of the proximal lobe, whereas the lower (IDA-facing) part of the proximal lobe likely contains the N-terminal regions of the “backbone subunits” DRC2 and DRC4.

**Fig. 5.**
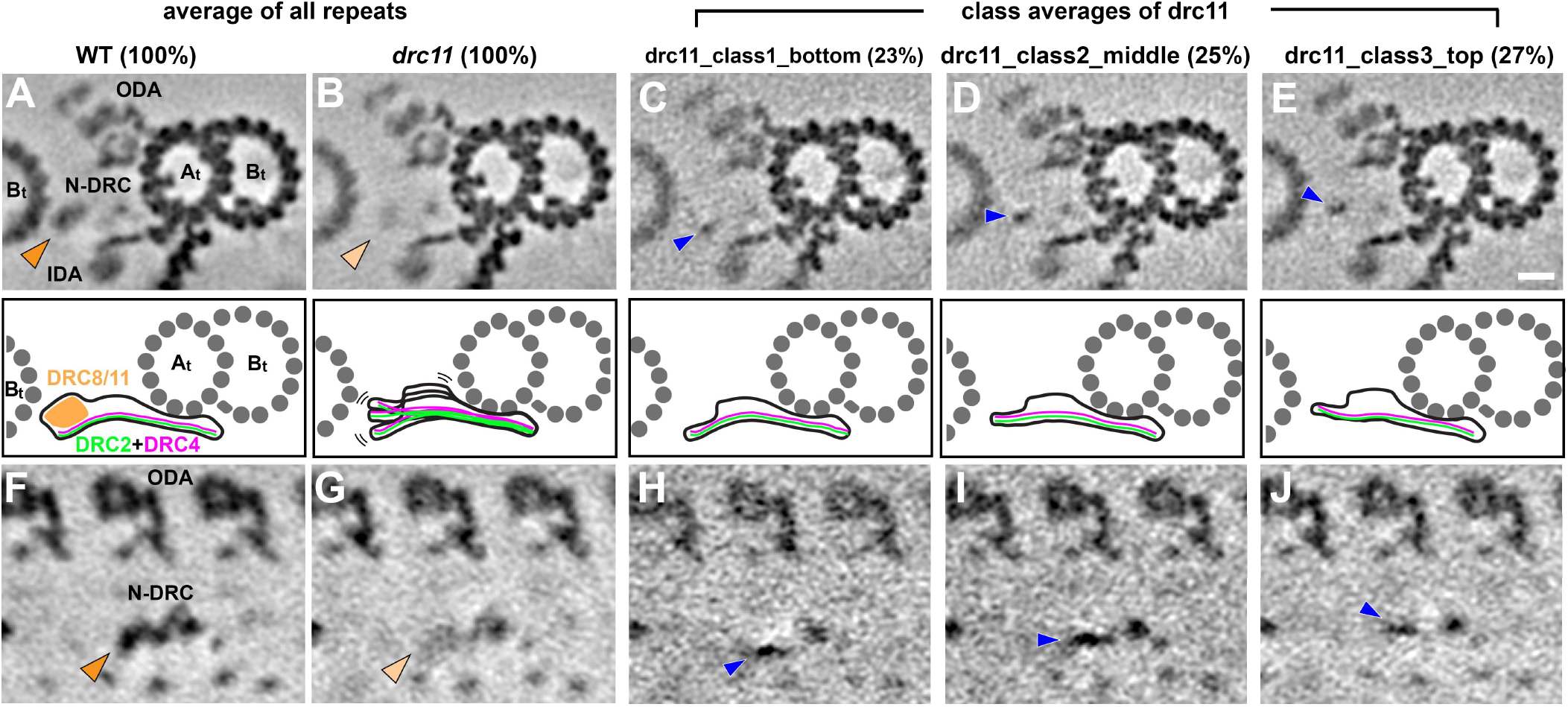
Classification analyses reveal that the DRC8/11 sub-complex is important for the stability/rigidity of the proximal lobe scaffold (DRC2/4). **(A-J)** Tomographic slices show the averaged axonemal repeats from wild-type (A and F) and *drc11* (B-E, and G-J) in cross-sectional (A-E) and longitudinal front (F-J) views. The first two columns show averages of all repeats (100%), whereas columns 3-5 show three classes of the *drc11* nexin-dynein regulatory complex (N-DRC) that vary in the positions of the proximal lobe (arrowheads). Note the relatively weak and blurry electron densities of the proximal lobe of the N-DRC in the average-all of *drc11* (light orange arrowheads in B and G). Classification analysis of this weak density in *drc11* revealed different positions occupied by the proximal lobe (blue arrowhead in classes 1-3; percentages of repeats are indicated for each class). Other labels: At, A-tubule; Bt, B-tubule; IDA, inner dynein arm; ODA, outer dynein arm. Scale bar: 10 nm.

### Rescue of *drc4* and *drc5* mutants with SNAP-tagged constructs clarifies the location of DRC4 and DRC5 in the N-DRC linker

Previous studies focusing on DRC subunits DRC1-6 showed that DRC1, DRC2, and DRC4 are coiled-coil proteins that extend from the inner junction between the A- and B-tubule through the N-DRC base plate and linker to the neighboring DMT (7, 14, 16) (Fig. 1*C*). Mutations in these proteins disrupt the assembly of multiple DRC subunits (12, 13, 22–25). In contrast, DRC3, DRC5, and DRC6 are located in the N-DRC linker region, and defects in DRC3, DRC5, or DRC6 have only small effects on the assembly of other DRC subunits or the overall structure of the N-DRC (7, 12, 13, 15, 16). Rescues of various *drc* mutants with tagged wild-type DRC genes allowed the localization of the N- and/or C-termini of DRC1 (N/C), DRC2 (N/C), DRC3 (N/C), DRC4 (C) and DRC5 (C) (Fig. 1*C*, *Right*). For example, the SNAP-tagged N- and C-termini of DRC3 have been unambiguously located in the L1-projection of the N-DRC linker that connects to IDA g (16). However, tagging the N-terminus of DRC4 with BCCP resulted in variable labeling throughout the proximal lobe. A C-terminal biotin carboxyl carrier protein (BCCP) tag of DRC5 appears to localize to the upper (ODA-facing) region of an unbranched part of the N-DRC linker and the electron density corresponding to the tag at the C-terminus of DRC5 was diffuse and blurred (7, 14). Moreover, an N-terminally BCCP tagged DRC5 construct was poorly expressed and could not be localized (14).

To address the uncertainty about the locations of the N-termini of DRC4 and DRC5, and the C-terminus of DRC5, we generated new SNAP- or BCCP-tagged constructs of *DRC4* and *DRC5* and used them to rescue the corresponding mutants *drc4 (pf2)* and *drc5 (sup-pf4)* (Fig. 6 and S3-S5). The forwarding swimming speeds of the rescued strains, i.e. *pf2-4;N-SNAP-DRC4, sup-pf4;DRC5-C-SNAP* and *sup-pf4;N-BCCP-DRC5*, were significantly faster than the mutants and increased to near wild-type levels (Fig. 6*B* and S5*B*). Western blots of purified axonemes demonstrated that the tagged proteins were expressed at endogenous levels and migrated at the sizes predicted for SNAP or BCCP tagged DRC subunits (Fig. 6*C*, 6*D* and S5*C*). The SNAP-DRC4 rescue also restored the assembly of other DRC subunits (Fig. 6*C*) that were previously shown to be missing in *pf2* mutant axonemes (13).

**Fig. 6.**
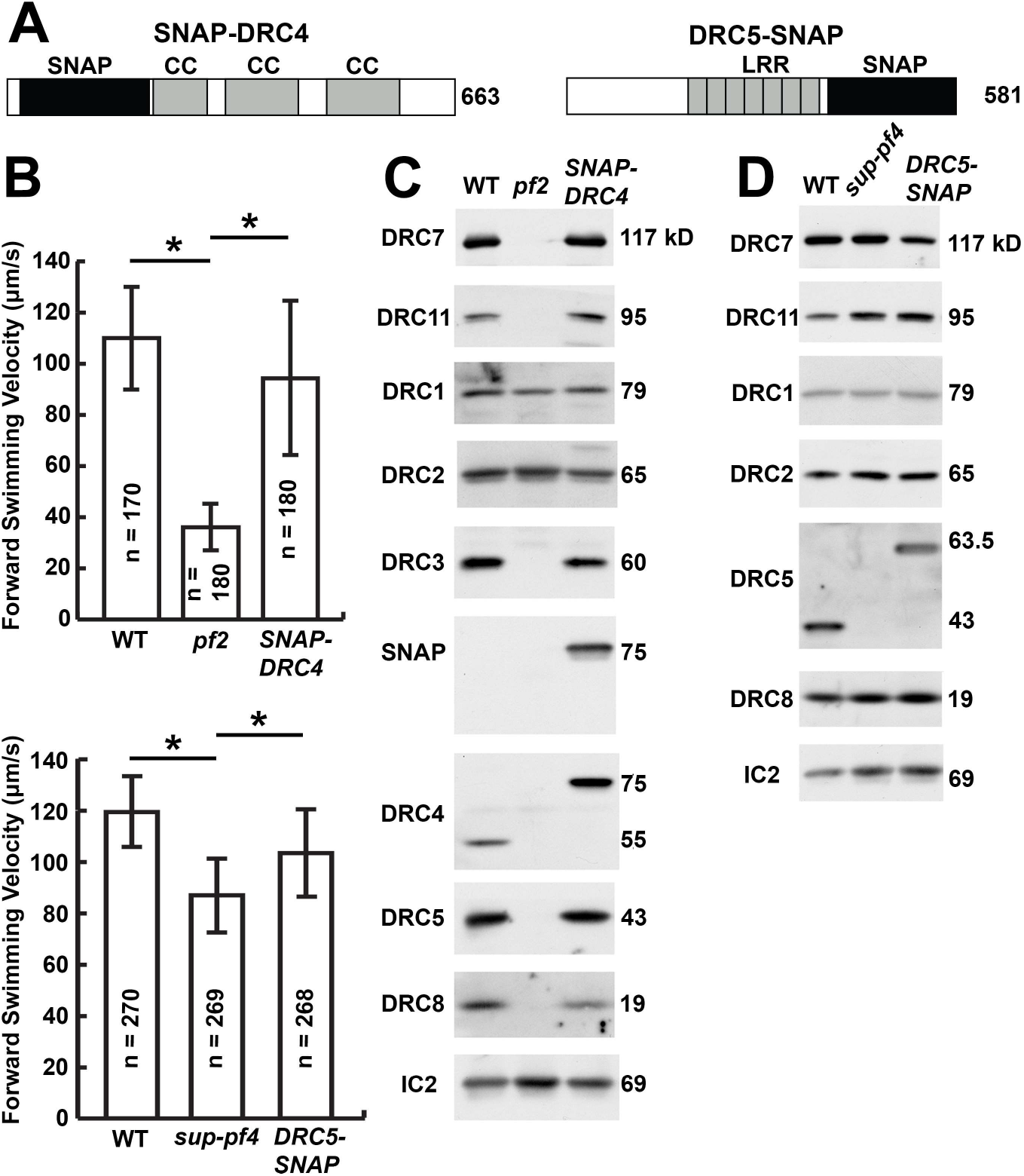
Rescue of *drc4* (*pf2*) and *drc5* (*sup-pf4*) mutants by transformation with SNAP-tagged DRC subunits. **(A)** The DRC4 and DRC5 protein sequences are drawn to scale, with the location of predicted polypeptide domains indicated: CC, coiled-coil domain; LRR, leucine-rich repeat; SNAP, SNAP tags introduced near the N-terminus of DRC4 and the C-terminus of DRC5. **(B)** The forward swimming velocities of *drc* mutants as measured by phase contrast microscopy are shown relative to those of wild type (WT) and rescued strains generated by transformation with the indicated SNAP-tagged *DRC* gene. The rescued strains were significantly faster (P < 0.05) than the *drc* mutants. **(C)** Western blots of axonemes isolated from WT, *drc* mutants, and SNAP-tagged rescued strains were probed with antibodies against several DRC subunits. Note the presence of bands that migrated at the sizes predicted for SNAP-tagged DRC subunits. An antibody against the IC2 subunit of the outer dynein arms served as a loading control.

Analyses of axonemes from the rescued strains by cryo-ET and sub-tomogram averaging revealed that the defects in N-DRC structure were completely rescued to WT structure and that the small SNAP and BCCP tags did not interfere with the proper assembly of the N-DRC or other axonemal structures (e.g. compare Fig. S6 *D-F* with *J-L*). After enhancing the visibility of the cloned tags in the rescued axonemes with streptavidin-nanogold labels, we were able to detect additional densities in the averages of the *SNAP-DRC4* and *DRC5-SNAP* axonemes (Fig. 7, S3 and S4), but not in the *BCCP-DRC5* averages (Fig. S5*J-L*). The failure to visualize the N-terminus of DRC5 is similar to previously reported results (Oda *et al* 2015). However, a clear density corresponding to the labeled N-terminus of DRC4 was found on the proximal side of the proximal lobe close to the neighboring B-tubule in *SNAP-DRC4* axonemes (Fig. 7*D-F* and S3). Taken together with the previously determined location of the C-terminus of DRC4, our data demonstrate that the DRC4 stretches along the entire length of the N-DRC, i.e. from its attachment site on the A-tubule near the inner DMT junction through the base plate to the proximal lobe. This is a similar structural arrangement to DRC1 and DRC2. The results reveal that DRC4 likely serves both as an anchor that attaches the N-DRC to the A-tubule and as a scaffold for the assembly of the other N-DRC subunits, similar to DRC1 and DRC2. Furthermore, averages of gold-labeled, *DRC5-SNAP* axonemes clearly showed the additional density of the gold nanoparticles that revealed the location of the DRC5 C-terminus at the center of the linker close, to where the proximal lobe emerges from the linker (Figs. *7G-I* and S4). Together with previous studies, our data provide a better understanding of the molecular organization of the N-DRC.

**Fig. 7.**
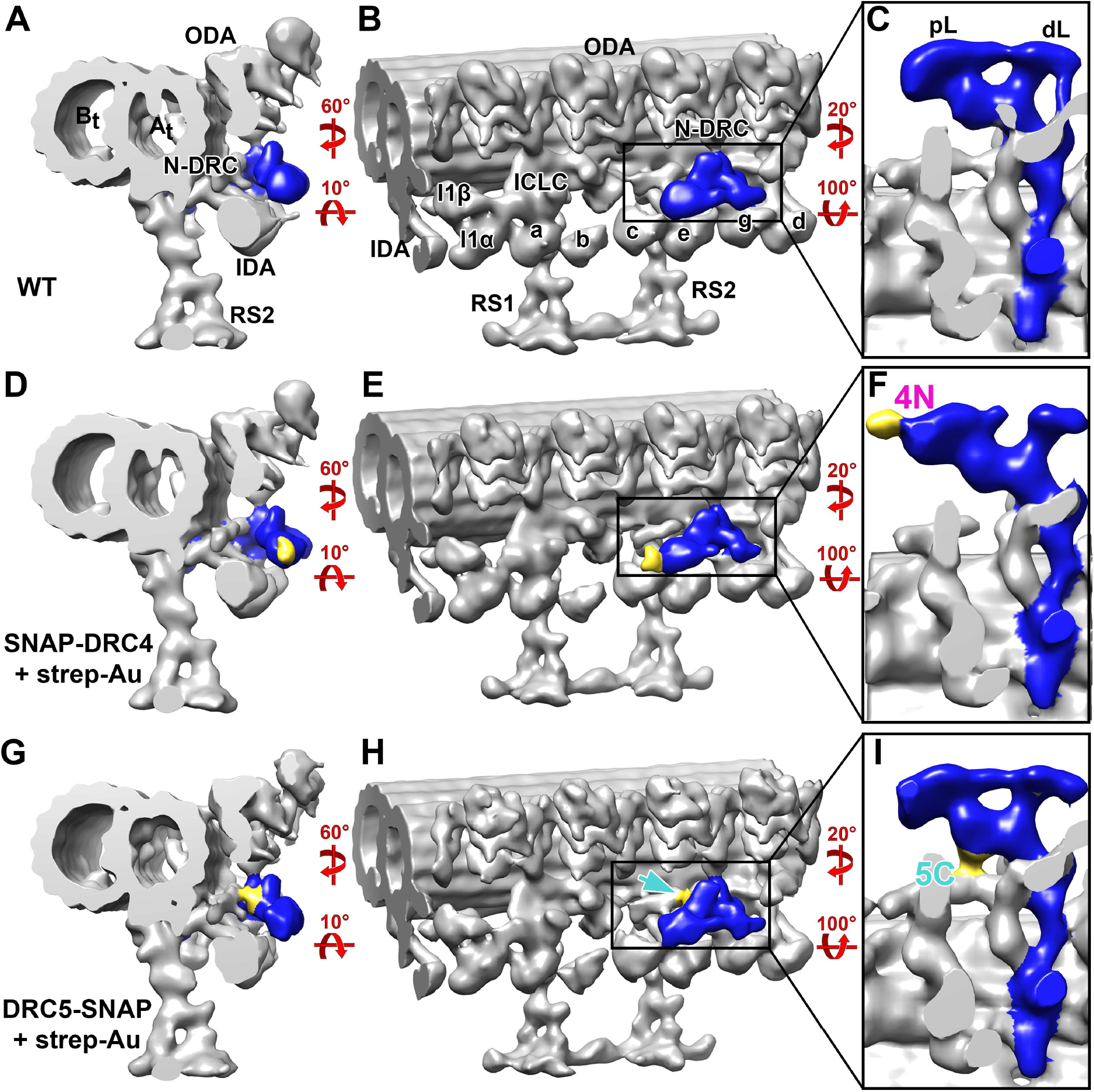
Precise localization of the N-terminus of DRC4, and the C-terminus of DRC5 by SNAP tag labeling and cryo-ET. Three-dimensional isosurface renderings show the structures of the 96-nm repeats in WT (A-C), labeled *SNAP-DRC4* (D-F), and labeled *DRC5-SNAP* (G-I) axonemes after cryo-ET and subtomogram averaging. Purified axonemes with SNAP-tagged DRC4 (D-F) or DRC5 (G-I) were labeled with streptavidin-Au (1.4 nm nanogold). Cross-sectional (A, D, and G), longitudinal front (B, E, and H) and enlarged longitudinal bottom views (C, F, and I; looking from the axoneme center outward) of the nexin-dynein regulatory complex (N-DRC, blue) show the additional density of the SNAP-biotin-strep-Au label (gold): at the proximal lobe (pL) of the N-DRC linker in the N-terminally labeled DRC4 axonemes (D-F), and at the center of the N-DRC linker in the C-terminally labeled DRC5 axonemes (G-I). Other labels: At, A-tubule; Bt, B-tubule; IDA, inner dynein arm; ODA, outer dynein arm; RS, radial spoke; dL, distal lobe; a–e and g, inner dynein arm isoforms; I1α/ I1β/ ICLC, α- and β-heavy chain, and intermediate-light chain complex of I1 dynein.

## DISCUSSION

Analysis by two-dimensional gel electrophoresis, immunoprecipitation, iTRAQ labeling and mass spectrometry has identified at least 11 N-DRC subunits (12, 13). In this study, by comparing the missing densities in the sub-tomogram averages of new *n-drc* mutants, we have revealed the location of three DRC subunits: DRC7 in the linker region including the OID-linker, and DRC8/11 in the upper portion of the proximal lobe. We also used gold labeling to determine the precise locations of the N-terminus of DRC4 and the C-terminus of DRC5 (Table 3). Combined with previous studies (7, 14, 16), we propose a revised model for the overall architecture of the N-DRC: The subunits DRC1, DRC2, & DRC4 form the core scaffold of the N-DRC and a platform for the assembly of functional subunits, including DRC3, DRC5, DRC6, DRC7, DRC8, and DRC11, with this backbone (Fig. 8 and Movie S4).

**Fig. 8.**
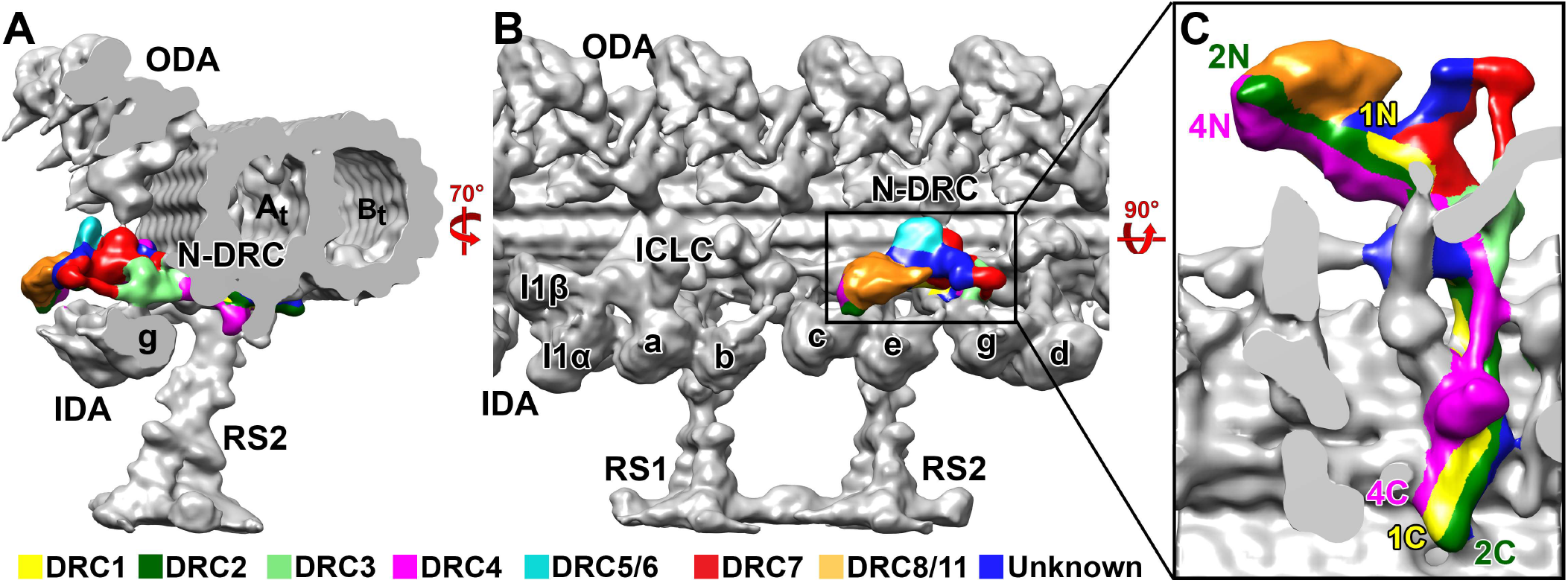
New model of the DRC subunit organization and N-DRC interactions with neighboring structures. The locations of the DRC subunits are summarized and colored in the wild type nexin-dynein regulatory complex (N-DRC) structure, which is shown in cross-sectional (A), longitudinal front (B) and enlarged bottom (C) views. The structural comparison between the averaged wild type axonemal repeat and deletion mutants of DRC subunits or mutant-rescues by tagged DRC subunits revealed the architecture of the N-DRC: the core scaffold extends along the entire length of the N-DRC and consists of DRC1 (yellow), DRC2 (dark green), and DRC4 (magenta); with this scaffold associate several functional subunits, i.e. DRC3 (light green), DRC5/6 subcomplex (cyan), DRC7 (red), and DRC8/11 subcomplex (orange). The composition of a few areas within the N-DRC remain unknown (dark blue). Other labels: At, A-tubule; Bt, B-tubule; IDA, inner dynein arm; ODA, outer dynein arm; RS, radial spoke; a–e and g, inner dynein arm isoforms; I1α/ I1β/ ICLC, α- and β-heavy chain, and intermediate-light chain complex of I1 dynein.

**Table 3.** Summary of polypeptides and structures present or absent in the strains used in this study.

| Components | Strains |  |  |  |  |  |  |  |
| --- | --- | --- | --- | --- | --- | --- | --- | --- |
|  | WT | *drc7* | *drc11* | *pf2* | *SNAP-DRC4* | *sup-pf-4* | *DRC5-SNAP* | *BCCP-DRC5* |
| **Polypeptides (kDa)** |  |  |  |  |  |  |  |  |
| DRC1 (79) | + | + | + | + | + | + | + | + |
| DRC2 (65) | + | + | + | + | + | + | + | + |
| DRC3 (60) | + | + | + | - | + | + | + | + |
| DRC4 (55) | + | + | + | - | + | + | + | + |
| DRC5 (43) | + | + | + | - | + | - | + | + |
| DRC6 (28) | + | + | + | - | + | - | + | + |
| DRC7 (177) | + | - | + | - | + | + | + | + |
| DRC8 (19) | + | + | - | - | + | + | + | + |
| DRC9 (46) | + | + | + | - | + | + | + | + |
| DRC10 (41) | + | + | + | - | + | + | + | + |
| DRC11 (95) | + | + | - | - | + | + | + | + |
| **Structure (kDa)** |  |  |  |  |  |  |  |  |
| Base Plate (~300) | + | + | + | +/- | + | + | + | + |
| Linker |  |  |  |  |  |  |  |  |
| Proximal lobe (~350) | + | + | - | - | + | + | + | + |
| Distal lobe (~100) | + | +/- | + | - | + | +/- | + | + |
| L2 protrusion (~200) | + | + | + | - | + | - | + | + |
| OID linker (~200) | + | - | + | - | + | + | + | + |

+ indicates that the component was present at wild-type levels; – indicates that the component was absent; +/- indicates that the component was present at reduced levels. Molecular mass of the N-DRC structure is estimated from the isosurface-rendering and indicated in parentheses (in kDa).

### DRC1, DRC2, and DRC4 form the backbone of the N-DRC

Cryo-ET and sub-tomogram averages of the axonemal repeat from *Chlamydomonas* null mutants of *drc1* (*pf3*), *drc2* (*ida6*), and *drc4* (*pf2*) previously revealed large-scale defects in the structure of the N-DRC, i.e. missing or reduced densities in both the base plate and the linker domains (7, 16, 23). Transformation with wild-type or tagged versions of DRC1, DRC2, or DRC4 fully restored the missing polypeptides and structures (7, 14, 16). The localization of the termini of DRC1, DRC2, and DRC4 in the previous and current studies suggests that all three subunits stretch along the entire N-DRC, i.e. from the inner junction to the neighboring DMT (14, 16). In addition, the use of advanced microscope hardware, such as a direct electron detector camera and Volta-Phase-Plate, has increased the resolution of our subtomogram averages, allowing us to visualize three filamentous structures in the base plate that are slightly twisted around each other like a rope (Fig. 8*C*). This is consistent with the high-resolution reconstruction of the wild-type *Tetrahymena* N-DRC using our recently developed TYGRESS software (26). These findings suggest that DRC1, DRC2, and DRC4 function as scaffold and play a pivotal role in the assembly of the entire N-DRC structure. Among the three backbone proteins, the sub-complex of DRC1 and DRC2 is the most critical as its absence in the *pf3* (*drc1*) and *ida6* (*drc2*) mutants caused reduced densities for the entire N-DRC base plate, including DRC4 (16, 23). Thus, DRC1 and DRC2 form likely the core filament at the very bottom of the N-DRC base plate, which directly associates with the A-tubule (Fig. 8*C*, yellow and dark green). DRC4 does not appear to be necessary for assembly of DRC1 and DRC2 into the base plate, as the *pf2* mutant has substantial remaining base-plate density, suggesting that DRC4 likely forms the filamentous structure on top of the DRC1/DRC2 sub-complex in the N-DRC base-plate (Fig. 8*C*, purple) (16, 23).

All scaffold components (DRC1, DRC2, and DRC4) are highly conserved in motile cilia and flagella, from unicellular algae *Chlamydomonas* to human. Genetic studies of the orthologues in different species have revealed that DRC1, DRC2, and DRC4 play essential and conserved roles in ciliary and flagellar motility. For example, an insertion into the *DRC4* locus in *Chlamydomonas* resulted in an altered ciliary waveform and slow swimming cells of the *pf2* strain (22). RNAi-mediated knockdown of the *DRC4* orthologue in trypanosomes revealed that DRC4 is required for flagellar motility and is essential for the viability of the bloodstream form of the parasite (27). The vertebrate orthologues of DRC4 are GAS11 in human and GAS8 in mice and zebrafish, and mutations in *GAS11* cause primary ciliary dyskinesia (28)

Previous cryo-ET studies failed to unambiguously identify the location of the N-terminus of DRC4. Oda et al. reported that the labeling density of the N-terminus of DRC4 appeared to be present in two locations, with one density at the proximal lobe and the other at the distal lobe. In contrast, in our studies the SNAP-tagged density clearly localized the N-terminus of DRC4 at the proximal lobe (Fig. 7*D-F*). The 96-nm averaged repeats of the *drc11* mutant showed a density that remains at the bottom (IDA-facing side) of the proximal lobe (Fig. 6). These results suggest that the N-terminal parts of DRC2 and DRC4 form the structural scaffold at the bottom of the proximal lobe, which can bind to the DRC8/11 subcomplex.

### DRC3, 5, 6, 7, 8, and 11 are attached to the N-DRC backbone and interact with other axonemal complexes

Loss of DRC3 (*drc3*), DRC5/DRC6 (*sup-pf4*), DRC7 (*drc7*), and DRC8/DRC11 (*drc11*) cause defects in certain regions of the N-DRC but have minimal effects on the assembly of other DRC subunits, unlike loss of the backbone subunits (7, 15, 16).

DRC5 is a leucine-rich repeat (LRR) protein that appears to be closely associated with DRC6 based on the absence of DRC6 in the DRC5 mutant *sup-pf4* (12, 29). Cryo-ET and subtomogram averaging revealed that the L2 protrusion of N-DRC linker, which is located in front of the OID-linker, is missing in *sup-pf4* axonemes (7) (Fig. S6*G* and *I*). The motility defects observed in the *Chlamydomonas sup-pf4* mutant are subtle. The *sup-pf4* mutant and the tubulin polyglutamylation (*tpg1*) mutant can both undergo reactivated motility in the presence of ATP, like WT cells, whereas the *sup-pf4;tpg1* double-mutant failed to reactivate motility in the presence of ATP. In addition, *sup-pf4;tpg1* axonemes were more prone to splaying, i.e. the nine DMTs separated more easily from each other, suggesting that DRC5/6 and B-tubule glutamylation of the neighboring DMT work together to maintain axonemal integrity and ciliary motility (30). However, it is not known which DRC subunit(s) make direct contact with the neighboring B-tubule. Here, cryo-ET localization of a tagged rescue strain suggested that the C-terminus of DRC5 is located in the center of the N-DRC linker domain, rather than close to the neighboring DMT. The N-terminal tag of BCCP-DRC5 could not be visualized (Fig. S5 *J-L*), even though Western blots of the rescued strain clearly showed that BCCP-DRC5 was expressed at WT levels (Fig. S5*C*), indicating that the addition of the BCCP tag does not interfere with the incorporation of DRC5 into N-DRC. Two possible reasons for failing to detect the location of the N-terminus of DRC5 might be that it is buried so deep in the structure of the N-DRC that the streptavidin-gold could not bind to the tag or that the N-terminal domain may be so flexible in position that the gold density would be averaged out in the sub-tomogram averages.

DRC7 is the largest N-DRC subunit (177 kDa) identified in *Chlamydomonas*. It contains a highly conserved transglutaminase-like (TGL) peptidase domain, which is predicted to bind tightly to glutamylated proteins, suggesting DRC7 may interact with glutamylated residues on the neighboring B-tubule. This idea is supported by our cryo-ET and sub-tomogram averaging images that show that a part of the distal lobe is missing in the *drc7* axonemes. However, the major defect in the *drc7* axonemes was seen in the middle-segment of the N-DRC linker, including a missing OID-linker that usually connects the N-DRC to the ODA row. The OID-linker is proposed to be part of the signal transduction pathway that regulates ciliary beating: signals are transferred from the CPC to the RS, then to the N-DRC and I1 dynein, finally reaching the ODAs via two routes; one OID-linker from the N-DRC and a second OID linker from the I1 intermediate-light chain complex, respectively (31–33). However, even though the defects in N-DRC structure seem to disrupt two important interfaces with neighboring structures, *drc7* mutant did not show significant defects in forward swimming velocity under standard culture conditions in *Chlamydomonas*. A possible explanation for the lack of a strong motility phenotype in *Chlamydomonas drc7* mutants might be that there are other – (partially) redundant – pathways for signal transduction. The N-DRC connects to the neighboring B-tubule via both the proximal and distal lobes, and the loss of the distal lobe in the *drc7* mutant might be compensated for by the remaining proximal lobe that still interacts with the neighboring B-tubule. In addition to the OID-linker from the N-DRC, also the I1 ICLC connects between the RS/IDA row to the ODAs, and these two signal transduction pathways between the IDA and ODA rows might be redundant for the activity control of the ODAs. Future study of double mutants in *Chlamydomonas* may reveal a more profound effect of the *drc7* mutation. Another possibility is that DRC7 might be critical for motility under more stringent conditions. In *Drosophila*, loss of the *DRC7* ortholog *CG34110* (*lost boys, lobo*) causes defects in sperm storage and fertility (34). Future studies will have to clarify the reasons for this more severe phenotype in flies, but a recent survey of motility genes expressed in *Drosophila* sperm has revealed that not all DRC subunits are expressed in the testis (35). Thus, it is also possible that loss of DRC7 may cause a more significant functional defects in other species.

DRC8 contains an EF-hand domain (a calcium-binding helix-loop-helix structural motif) that could bind Ca^2+^, and DRC11 contains a calmodulin-binding motif, suggesting these two subunits may contribute to the calcium regulation of flagellar motility. DRC11 also contains an “ATPases Associated with diverse cellular Activities” (AAA)-domain that could mediate nucleotide-sensitive conformational changes. Our studies of the *drc11* mutant showed that DRC11 is required for the assembly of DRC8, and DRC8/11 probably form a sub-complex localized to the proximal lobe of the linker domain that contacts the neighboring B-tubule (Fig. 4). Thus, we propose that the DRC8/11 subunits regulate cilia motility by modulating contacts with the neighboring B-tubule, which could be important for transforming interdoublet sliding to ciliary bending, and/or provide signal feedback. The ortholog CMF22 in *Trypanosoma brucei* is required for normal flagellar beating patterns and propulsive cell motility (36). However, in *Chlamydomonas*, we only observed small decreases in the swimming velocity of the *drc11* mutant compared to wild-type cells (Fig. 2*B*). One possible explanation may be that the N-DRC connects to the neighboring B-tubule via both the proximal lobe and the distal lobe. The proximal lobe is missing in the *drc11* mutant, but the remaining distal lobe could interact with the B-tubule and transform the interdoublet sliding to bending. It is interesting that loss of the DRC8/11 subcomplex leads to positional flexibility of the remaining N-DRC between the IDA and ODA row, because in a recent study of actively beating sea urchin flagella we found that the N-DRC adopted three different conformational states (Lou et al. in preparation). Structurally, these states varied in their position between the IDA and ODA rows and showed a bend-direction specific distribution in the beating flagella, suggesting that the conformations were related to the different activity state of the dyneins (Lou et al. in preparation).

Orthologues of N-DRC subunits have been found in motile cilia, both “9 + 2” and “9 + 0”, of many species (13, 37, 38), but not in non-motile primary “9 + 0” cilia (39, 40). The fact that the N-DRC seems to be present in all motile cilia, even in motile “9 + 0” cilia that can lack almost any other major axonemal complexes, such as the CPC, RSs, and IDAs or ODAs (6, 37), suggests an essential and central role for the N-DRC in regulating ciliary motility. Our results provide insights into several important aspects of N-DRC assembly and function, from revealing the precise location of several DRC subunits, the overall organization of the N-DRC with a three subunits core-complex (DRC1/2/4) that serves as the scaffold for the assembly of the “functional subunits” (DRC3/5-8/11), shedding light on the molecular mechanism by which the N-DRC regulates dynein activity and thus flagellar beating.

## METHODS

### Culture conditions, strain construction, and identification of new mutations

Strains, including wild-type *Chlamydomonas reinhardtii* and CLiP mutants (Table S1), and mutant strains rescued with tagged DRC subunits were cultured as previously described (13) (see supplemental information for more details on the identification and characterization of *drc* mutants and the construction of epitope-tagged *N-DRC* strains). The *Chlamydomonas* strain *cw15* that was used as the background strain for the CLiP library, is cell wall-less but assembles flagella resembling those of wild type (*cc-125*).

### Phase contrast, fluorescence microscopy and measurements of swimming velocity

Transformants and candidate *drc* mutants were screened by phase contrast microscopy using a 20x or 40x objective, a halogen light source, and a red filter on a Zeiss Axioskop. Forward swimming velocities were measured using a Rolera-MGi EM-CCD camera (Q Imaging, Tucson, AZ) and the Metamorph software package, version 7.6.5.0 (Molecular Devices, Downington, PA) as previously described (13). The forward swimming velocities of the CLiP mutants and CLiP-mutant rescues were compared with the “wild-type” like CLiP background strain *cw15*, whereas *pf2, suppf4, SNAP-DRC4, DRC5-SNAP, BCCP-SNAP* were compared to the wild type strain *cc-125*. Some transformants were screened for the presence of the SNAP or BCCP tag by immunofluorescence microscopy (41).

### Preparation of purified axonemes for SDS-PAGE, Western Blotting, iTRAQ labeling and mass spectrometry

*Chlamydomonas* isolated axonemes were prepared as previously described (13, 42). Briefly, Chlamydomonas cells were collected by centrifugation and resuspended in pH shock buffer containing 10 mM HEPES, pH 7.4, 1 mM SrCl2, 4% sucrose, and 1 mM dithiothreitol (DTT). Flagella were detached from cells by adding 0.5 M acetic acid to reduce the pH to 4.3. After 80 s, the pH was increased to 7.4 by adding 1 M KOH. The flagella pellets were then demembranated by adding 1% Igepal CA 630 (Sigma-Aldrich) to the solution with gentle rotation for 20 min at 4°C. Axonemes were collected by centrifugation at 10,000 × g for 10 min and resuspended in HMEEN (10 mM HEPES, 5 mM MgSO_4_, 1 mM EGTA, 0.1 mM EDTA, 30 mM NaCl, pH 7.4) plus 1 mM DTT and 0.1 μg/ml protease inhibitors (leupeptin, aprotinin, pepstatin). Samples were separated on 5-15% polyacrylamide gradient gels, transferred to Immobilon P, and probed with the different antibodies listed in Table S3 as described by (13).

For iTRAQ labeling and mass spectrometry, isolated axonemes were washed with 10 mM Hepes pH 7.4 to remove salt, DTT, and protease inhibitors, then resuspended in 0.5 M triethylammonium bicarbonate pH 8.5 and processed for trypsin digestion and iTRAQ labeling as described in (13, 41). Samples were combined and fractionated offline using high pH, C18 reversed phase chromatography. Column fractions were vacuumed dried, resuspended in solvent (98:2:0.01, water:acetonitrile:formic acid), and loaded in 1-1.5 μg aliquots for capillary LC using a C18 column at low pH. The C18 column was mounted in a nanospray source directly in line with a Velos Orbitrap mass spectrometer (Thermo Fisher Scientific, Inc., Waltham, MA). Online capillary LC, MS/MS, database searching, and protein identification were performed as previously described (13, 43) using ProteinPilot software version 5.0 (AB Sciex, Foster City, CA) and the most recent version (v5.5) of the *Chlamydomonas* database (https://phytozome.jgi.doe.gov/pz/portal.html). The bias factors for all samples were normalized to alpha and beta-tubulin.

Wild-type (*cw15*) and *drc* axonemes were each labeled with two different iTRAQ reagents in a 4-plex experiment (technical replicates), and then each 4-plex experiment was repeated with new samples (biological replicates). For the *drc7* (197909) mutant, 1166 proteins were detected with high confidence at a false discovery rate of 5% in the first 4-plex experiment, and 701 proteins were detected in the second experiment. For the *drc11* (068819) mutant, 949 proteins were detected in the first 4-plex experiment, and 768 proteins were detected in the second experiment. The datasets were further filtered to identify those proteins who were identified by at least 5 peptides and whose mutant/WT ratios were significantly different (P value <0.05) from WT/WT in all replicates.

### Axoneme preparation and SNAP-gold labling for cryo-ET

*Chlamydomonas* axonemes for cryo-ET were prepared as mentioned above. The only difference is that the final axoneme pellets were resuspended in HMEEK buffer (30 mM HEPES, 5 mM MgSO4, 1 mM EGTA, 1 mM EDTA, and 25 mM KCl, pH 7.4,). 3 μl of the isolated axonemes was added to glow-discharged R2/2 holey carbon-coated grids and gently mixed with 1 μl of 10-times-concentrated, BSA-coated 10-nm gold solution (44). After ~2 seconds of backside blotting with filter paper, the grid was rapidly plunge frozen in liquid ethane with a homemade plunge freezer. Frozen grids were stored in liquid nitrogen until used.

For SNAP-gold labeling samples, the SNAP tag was labeled with gold as previously described (16). Briefly, 1 μl of 1 mM BG-biotin or BG-(PEG)12-biotin (New England Biolabs; PEG linker available on request) was added to 200 μl of the purified axonemes in HMEEK buffer and incubated overnight at 4 °C. Unbound BG-substrate was removed by centrifugation at 10,000 g for 10 min at 4 °C and three wash steps with HMEEK buffer. The axoneme pellet was resuspended in 200 μl of HMEEK buffer. After adding 5 μl of 80 μg/ml 1.4-nm-sized streptavidin nanogold particles (strep-Au, Nanoprobes, Inc.), the suspension was incubated at 4 °C for 4 h. After adding 800 μl of HMEEK buffer, the labeled axonemes were pelleted by centrifugation at 10,000 g for 10 min at 4 °C and resuspended in 200 μl of HMEEK buffer.

### Cryo-Electron Tomography

Vitrified grids were imaged using a Titan Krios transmission electron microscope (used for all samples unless otherwise noted) or Tecnai F30 (used for wild-type, *SNAP-DRC4, DRC5-SNAP, BCCP-DRC5* axonemes) both operated at 300 kV. Images were captured using a 4k × 4k K2 direct detection camera (Gatan) on the Titan Krios at a magnification of 26,000x (yielding a scaling of 5.5 Å per pixel) or a 2k × 2k charge-coupled device camera (Gatan, Pleasanton, CA) on the Tecnai F30 at a magnification of 13,500x (yielding a scaling of 10.0 Å per pixel). Tilt series were collected from 60° to −60° in 2° steps using dose-symmetric method on the Titan Krios. Counting mode of the K2 camera was used and for each tilt image 15 frames (0.4 s exposure time for each frame) were recorded. Both cameras were placed behind a post-column energy filter (Gatan) that was operated in zero-loss mode (20-eV slit width). Data acquisition was performed using the microscope control software SerialEM (45) in low-dose mode, and the total electron dose per tilt series was limited to ~100 e/Å^2^.

### Image Processing

The frames of each tilt series image collected on a K2 camera were aligned and then merged using the script extracted from the IMOD software package (46) to generate the final tilt serial data set. Tomograms were reconstructed using fiducial alignment and the back-projection method in the IMOD package (46). Subtomogram averaging was performed using the PEET software (19, 20) to average the 96-nm axonemal repeat units from 3D reconstructed axonemes. The resolution of the DMT averages was estimated using the 0.5 criterion of the Fourier shell correlation method (Table S2). The Chimera package (47) was used for 3D visualization and surface rendering. Mass estimations of the DRC complex and sub-volumes were calculated using the average density of 1.43 g/cm^3^ for proteins (48) and after normalizing the isosurface-rendering threshold to the mass of microtubules.

## Acknowledgments

We thank Chen Xu (Brandeis University), Zhenguo Chen and Daniel Stoddard (UT Southwestern Medical Center) for management of the electron microscope facilities and training. The UT Southwestern Cryo-Electron Microscopy Facility is supported in part by the CPRIT Core Facility Support Award RP170644. We also thank Gang Fu for assisting with tomogram reconstruction. We are grateful to Cai Kai for critical reading of the manuscript. We also thank LeeAnn Higgins and Todd Markowski in the Center for Mass Spectrometry and Proteomics (CMSP) at the University of Minnesota for assistance with mass spectrometry of the iTRAQ labeled samples. This center is supported by multiple grants including National Science Foundation (NSF) Major Research Instrumentation grants 9871237 and NSF-DBI-0215759 as described at www.cbs.umn.edu/msp/about. We also acknowledge Matt Laudon and the Chlamydomonas Genetics Center (University of Minnesota) for strains. This facility is supported by the National Science Foundation Living Stock Collections for Biological Research program grants 0951671 and 00017383. The antibody to RSP16 was generously provided by Pinfen Yang (Marquette University) and the antibody to DRC1 by Win Sale (Emory University). This study was funded by the following grants: National Institutes of Health R01GM083122 to D.N. and R01GM055667 to M.E.P.

## Data deposition

The three-dimensional averaged structures of 96-nm axonemal repeats have been deposited in the Electron Microscopy Data Bank (EMDB) under accession numbers EMD-20338, EMD-20339, EMD-20340 and EMD-20341.

## Author Contribution

D.N. and M.E.P. designed research. L.G., K.S., S.Y., A.D., M.G., and T.N. carried out axoneme preparation, cryo-electron tomography and sub-tomogram averaging. D.T. generated genomic and cDNA constructs, characterized insertion sites in new *drc* mutants, transformed and screened mutant strains for rescue. R.B. isolated axonemes and performed Western blot and iTRAQ analyses. K.A. and J.S. assisted with the screening of transformants and performed all measurements of swimming velocities. L.G., K.S., D.T., R.B., and K.A. contributed to data analysis and interpretation. L.G., M.E.P., and D.N. wrote the manuscript with input from all authors.

## SI Materials and Methods

### Culture conditions, strain construction, and identification of new mutations

Cells were maintained on Tris-acetate phosphate (TAP) medium but occasionally cultured overnight in minimal liquid medium for measurements of swimming velocity or axoneme preparation.

### SNAP tagging the N-terminus of DRC4

To generate the *SNAP-DRC4* strain, a genomic fragment encoding the N-terminal region of *DRC4* was amplified using the PCR primers listed in Table S4 and subcloned into pGEM T-easy (Promega Corp, Madison, WI) to generate subclone *pf2-DT6-7*. The *pf2-DT6-7* subclone contains a unique *NruI* site at the start of exon 2 and *MfeI* and SgrA1 sites on either side of exon 2. A *Chlamydomonas* version of the SNAP tag (1) was amplified using PCR primers containing *NruI* restriction sites (Table S4) and cloned into the *NruI* site of the pf2DT6-7 subclone using InFusion cloning (Takara Bio USA, Mountain View, CA). SNAP positive subclones were identified by PCR and sequenced to confirm the reading frame and sequence. This construct, pf2-DT6-7 SNAP, encodes a new alanine residue at amino acid 19 followed by the SNAP tag at amino acids 20-210 (Table S3). The pf2-DT6-7 SNAP subclone was digested with *MfeI* and SgrA1, gel purified and ligated into the original *pf2* genomic clone (p9B11-X1.2)(2). The sequence of the final pf2-N-SNAP construct was verified, and then the clone was digested with *AatII* and cotransformed with the selectable marker pSI103 (encoding the *AphVIII* resistance gene) into the *pf2-4* strain. Transformants were selected by growth on TAP plus 10 μg/ml paromomycin, and 212 colonies were screened for the rescue of motility by phase contrast microscopy (15 rescues) and the presence of the SNAP tag by immunofluorescence microscopy (14 positive). Positive transformants were rescreened on Western blots of whole cell extracts. Three single colony isolates were characterized by measurements of forwarding swimming velocity and Western blots of axonemes. Isolate 13-1 was saved for further study.

### SNAP tagging the C-terminus of DRC5

To generate the *DRC5-SNAP* strain, the SNAP tag was amplified using PCR primers containing *FseI* sites (Table S4) and subcloned into a 7.4 kb subclone of the *DRC5* gene containing a mutated stop site(3). SNAP positive subclones were identified by PCR and sequenced to verify the reading frame and sequence. This construct, *DRC5-SNAP*, adds three amino acids (WPA) at the end of the DRC5 sequence followed by 194 amino acids of the SNAP tag (Table S3). The DRC5-C-SNAP construct was linearized with *KpnI* and co-transformed with pSI103 into *sup-pf4* as described above. Thirty-two transformants were screened for the presence of the SNAP tag on Western blots of cell extracts, and three SNAP positive strains were identified. Following single colony isolation, the three strains were analyzed by phase contrast microscopy and Western blots of axonemes. Isolate 2D and 4D are saved for further study.

### BCCP tagging the N-terminus of DRC5

To insert a tag near the amino terminus of DRC5, the DRC5 plasmid was digested with *EcoRV* and gel purified. Then a BCCP tag was amplified using the PCR primers shown in Table S4 and inserted into the *EcoRV* digested plasmid using In-Fusion cloning. The BCCP-DRC5 construct adds 90 amino acids after amino acid 42. The construct was linearized with *BamHI* and co-transformed with pSI103 into *sup-pf4* as described above, and 192 transformants were screened for presence of BCCP tag by immunofluorescence of fixed cells (4/50 positive) and Western blots of axonemes (1/4 positive). Isolate 5-2 was saved for further study.

### Characterization of *drc7* mutations

Ten potential *drc7* mutations were identified in the CLiP library of insertional mutants(4, 5), but only four strains were reported to contain high confidence (95%) insertions in introns or at an intron/exon boundary. Mutations in two strains, LMJ.RY0402.080<u>526</u> and LMJ.RY0404.197<u>909</u>, were confirmed by genomic PCR (Table S4 and Fig. S1*A-C*). The 080<u>526</u> insertion appeared to result in alternative splicing of the *DRC7* transcript and the assembly of two DRC7 related bands in isolated axonemes (Fig. S1*D*). The 197<u>909</u> insertion appeared to be a null allele (Fig. 2) and was saved for further study.

### Characterization of *drc11* mutations

Eleven potential *drc11* mutations were identified in the CLiP library, but only five strains were reported to contain high confidence insertions, three in the 3’UTR and two in introns. The two intron mutations (130<u>937</u> and 068<u>819</u>) were confirmed by genomic PCR (Table S4 and Fig. S1*E-G*) and both disrupted assembly of DRC11 (Fig. 2 and S2*C*).

To rescue the *drc11* strains, a genomic BAC clone (39g16) was obtained from Clemson University (Clemson, South Carolina) and a cDNA clone (#AV628460) from the Kazusa DNA Research Institute (Kisarazu, Japan). The *DRC11* cDNA clone was amplified as two separate PCR products using the primers listed in Table S4 for subcloning into a modified version of the pIC2-MCS-C-BCCP-AphVIII plasmid(6). This plasmid was modified in three steps: first, the BCCP tag was replaced with a SNAP tag, then *DRC11* was cloned into the multiple cloning site (MCS), and finally, *AphVIII* was replaced with *AphVII*. Step 1: The plasmid was digested with EcoRI and EcoRV to release BCCP. The SNAP tag was amplified with primers containing an EcoRI site on one side and EcoRV site on the other (Table S4) and ligated into the digested plasmid using InFusion (Takara Bio USA) to generate pIC2-MCS-C-SNAP-AphVIII. Step 2: The plasmid was digested with BamHI and EcoR1, and the two PCR products encoding DRC11 were inserted into the MCS using the NEBuilder HiFi DNA assembly kit (New England Biolabs, Ipswich, MA) to generate pIC2-DRC11-C-SNAP-AphVIII. Step 3: The plasmid was digested with *HindIII* to release the *AphVIII* gene (encoding resistance to paromomycin). The *AphVII* gene (encoding resistance to hygromycin) was amplified from the pHyg3 plasmid using the primers listed in Table S4 and incorporated into the final plasmid using the NEBuilder HiFi DNA assembly kit (7). The orientation and reading frame of each plasmid was confirmed by DNA sequencing. The final plasmid (pIC2-DRC11-C-SNAP-AphVII) encodes the first exon of LC2 followed by DRC11-C-SNAP (Table S3). This plasmid was digested with *SpeI* and transformed into the *drc11-819* strain. Positive transformants were selected on TAP plus hygromycin B (Sigma-Aldrich, St. Louis, MO), and 58 colonies were screened for expression of DRC11-SNAP on Western blots of cell extracts. One SNAP positive clone was single colony isolated and saved for further study.

**Fig. S1.**
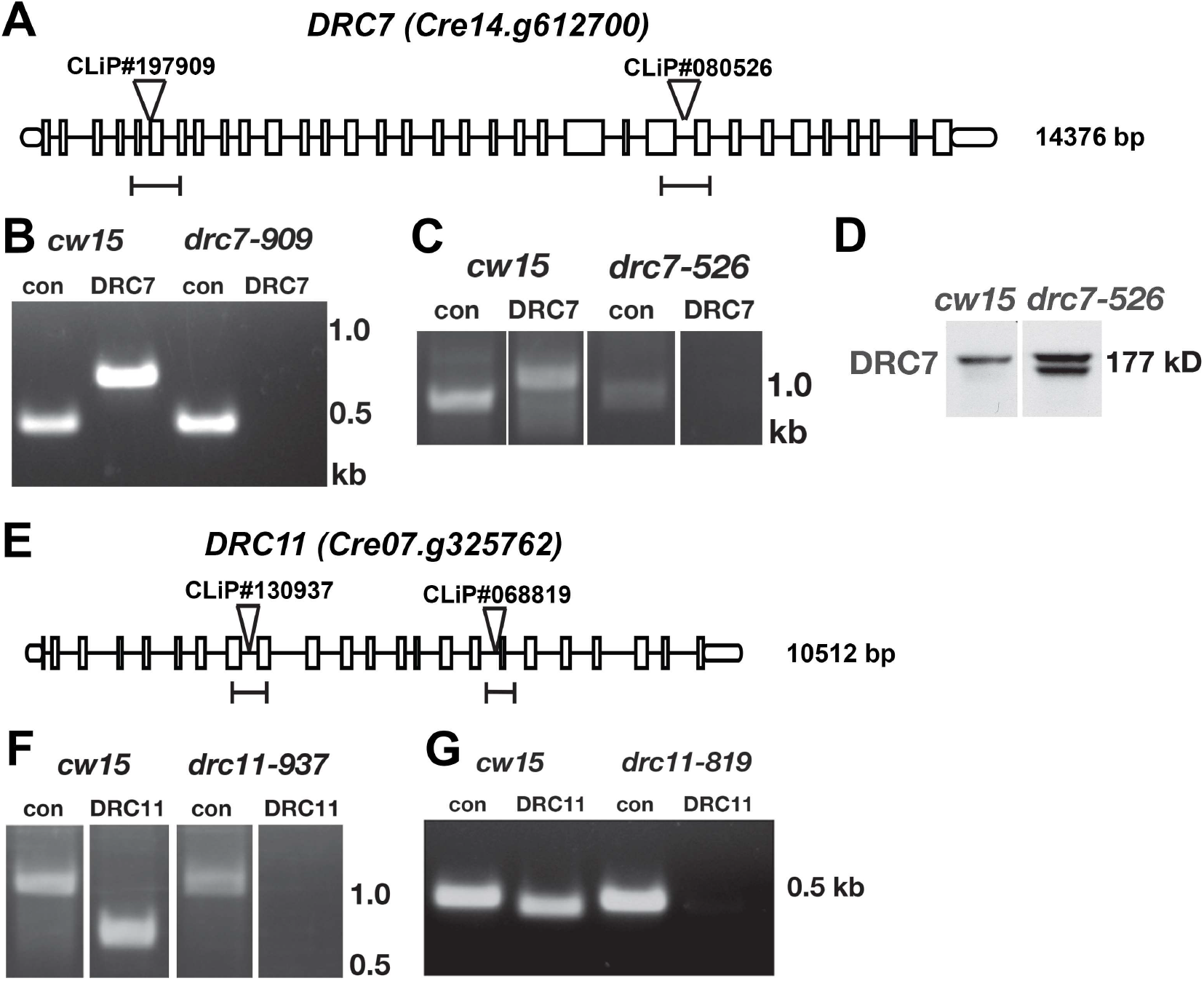
**(A, E)** The *DRC7* (*Cre14.g612700*) and *DRC11* (*Cre07.g325762*) genes are shown diagrammatically, with the 5’ and 3’UTRs as circular ellipses, exons as boxes, and the intervening introns as lines. The approximate locations of the plasmid insertion sites in the two mutant strains chosen for more detailed study are indicated by the inverted triangles. The sizes and locations of the PCR products in the wild-type genes are indicated by the lines below. **(B, C, F, G)** Agarose gel electrophoresis of the PCR products obtained with control primers (con) and the *DRC7* or *DRC11* primers listed in Table S4 using DNA from the background strain (*cw15*) or the mutant strains *drc7-909* (CLiP #197909), *drc7-536* (CLiP #080526), *drc11-937* (CLiP #130937), or *drc11-819* (CLiP #068819). **(D)** Western blots of axonemes indicated that the intron mutation in the *drc7-526* strain (CLiP #080526) results in alternative splicing and assembly of two DRC7 related polypeptides, one similar in size to the WT protein and one slightly truncated protein.

**Fig. S2.**
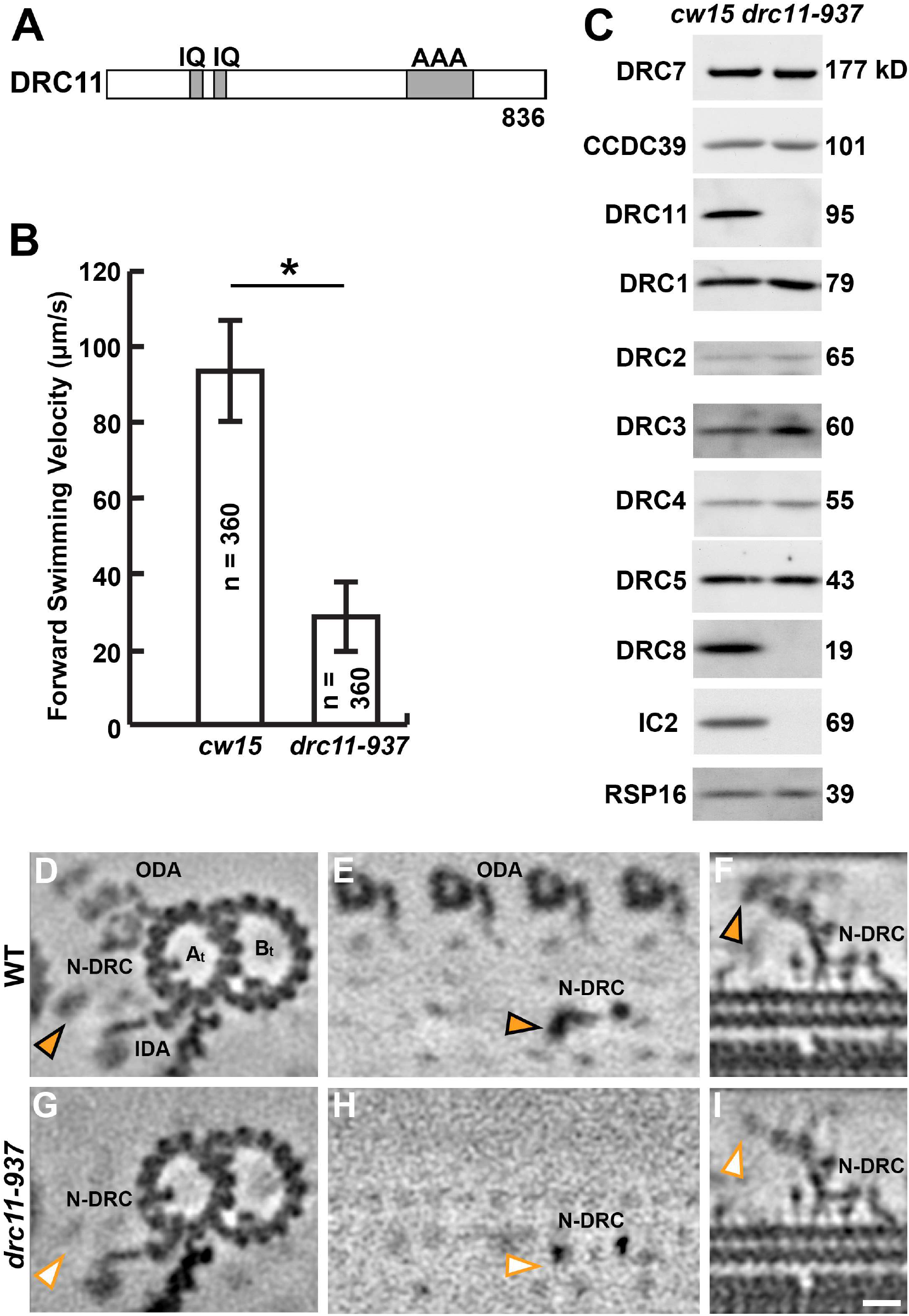
The *drc11-937* strain (CLiP #130937) not only lacks DRC11 and DRC8, which causes structural defects in the proximal lobe of the N-DRC linker, but also lacks the outer dynein arms. **(A)** Diagram of the DRC11 polypeptide sequence with the predicted domains indicated. IQ, calmodulin binding domain; AAA, <u>A</u>TPase <u>a</u>ssociated with diverse cellular activities domain. **(B)** The forward swimming velocities of the background strain (*cw15*) and the *drc11-937* strain (CLiP #130937) were measured by phase contrast microscopy. The total number of cells measured for each strain (n) is indicated. The *drc11-937* strain was significantly slower than the control *cw15* strain (p < 0.05). However, we were unable to rescue the motility defects by co-transformation with a BAC clone (39g16) containing the WT *DRC11* gene (0 positives out of 538 transformants screened) because of a second, unmapped mutation that disrupted the assembly of the ODAs. **(C)** Western blots of isolated *drc11-937* axonemes were probed with antibodies against several DRC subunits. Antibodies against CCDC39 and RSP16 served as loading controls. Antibodies against the IC2 subunit (an intermediate chain of ODA) revealed a defect in ODA assembly. **(D-I)** Tomographic slices of the averaged 96-nm axonemal repeats from *Chlamydomonas* wild type (D-F) and *drc11-937* (G-I) viewed in cross-sectional **(D, G)** and longitudinal views **(E, F, H, I).** Electron densities of the proximal lobe of in the nexin-dynein regulatory complex (N-DRC) were observed in wild type (orange arrowheads), but were missing in *drc11-937* (white arrowheads). Note the *drc11-937* mutant is also missing the outer dynein arms (ODA). Other labels: At, A-tubule; Bt, B-tubule; IDA, inner dynein arm. Scale bar: 10 nm.

**Fig. S3:**
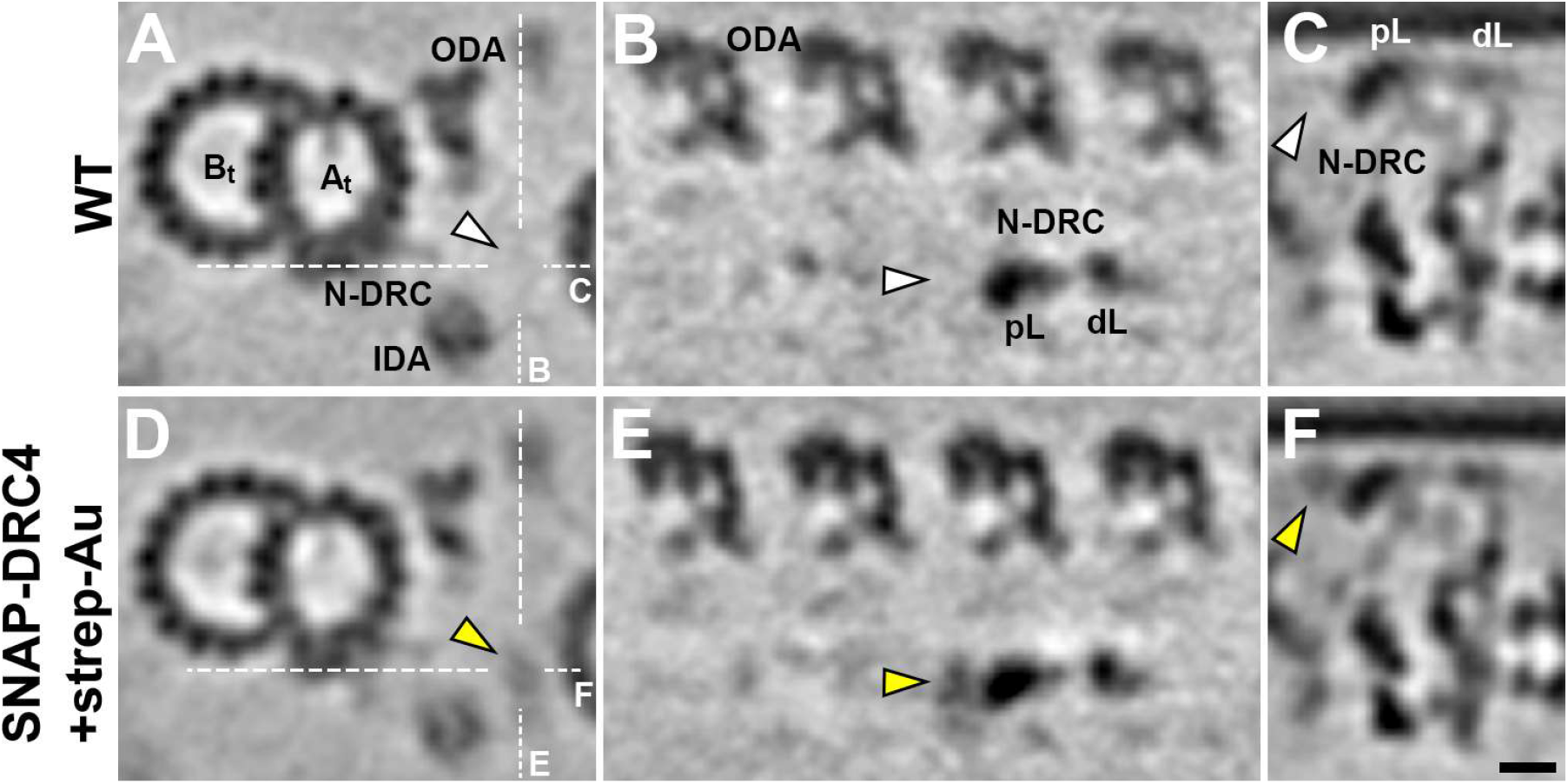
Comparison between the averaged 96-nm axonemal repeats from *Chlamydomonas* WT and the *pf2* mutant that was rescued with SNAP-DRC4. **(A-F)** Tomographic slices of the averaged 96-nm axonemal repeats from wild type (A-C) and *pf2-4;N-SNAP-DRC4* (D-F) viewed in cross-sectional (A, D) and longitudinal views (B, C, E, F). White lines indicate the locations of the slices in the respective panels. Note the additional electron density (yellow arrowheads in D-F) corresponding to the streptavidinnanogold marking the location of the N-terminus of DRC4 at the proximal side of the proximal lobe (pL) of the nexin-dynein regulatory complex (N-DRC). Other labels: At, A-tubule; Bt, B-tubule; dL, distal lobe; IDA, inner dynein arm; ODA, outer dynein arm. Scale bar: 10 nm.

**Fig. S4.**
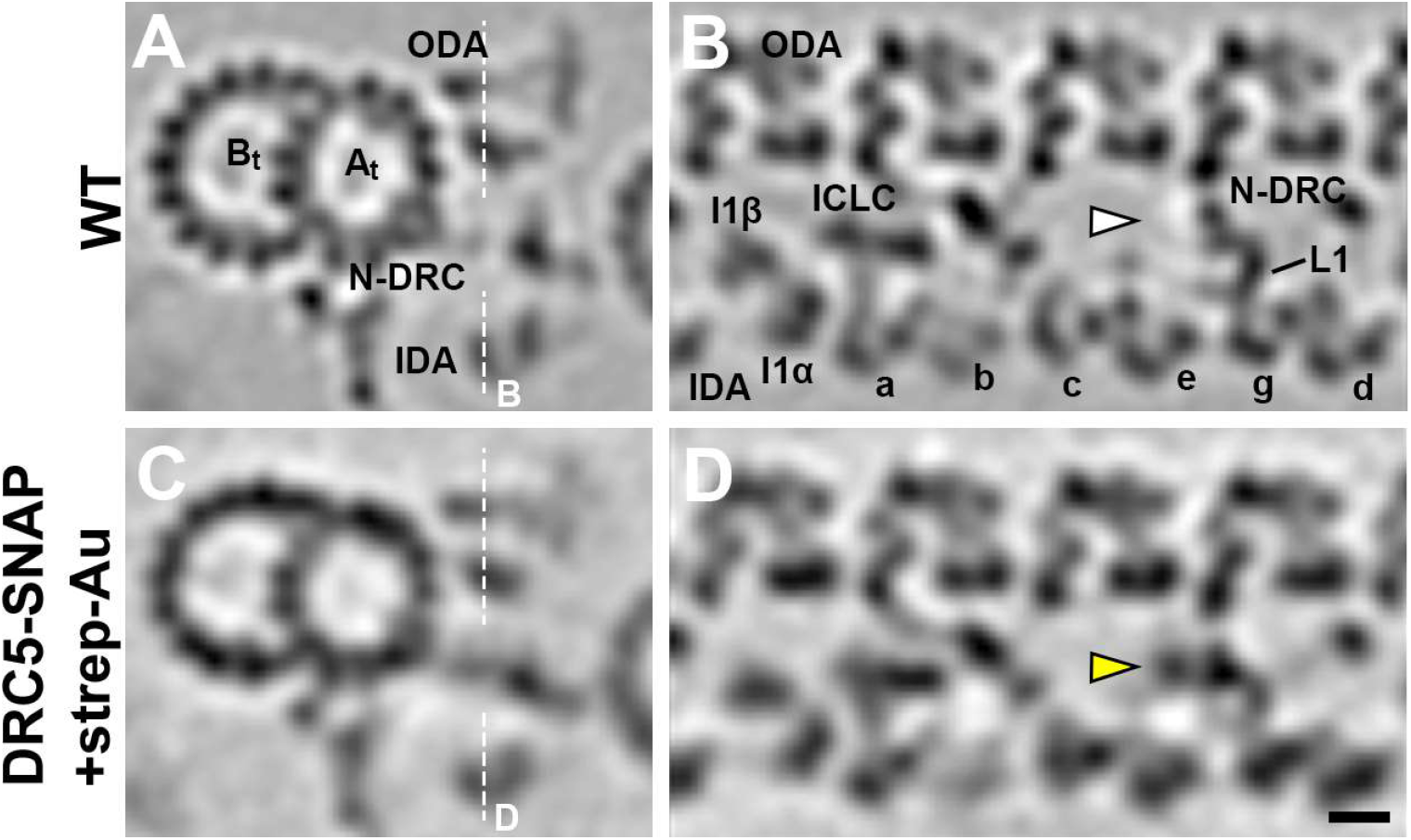
Localization of the SNAP-tagged C-terminus of DRC5 in *Chlamydomonas* axonemes. **(A-D)** Tomographic slices of the averaged 96-nm axonemal repeats from wild type (A and B) and *sup-pf4;DRC5-C-SNAP* (C and D) viewed in cross-sectional (A and C) and longitudinal views (B and D). White lines indicate the locations of the slices in the respective panels. Note the additional electron density (yellow arrowheads in D) corresponding to the streptavidin-nanogold marking the location of the C-terminus of DRC5 near the center of the nexin-dynein regulatory complex (N-DRC) linker. Other labels: At, A-tubule; Bt, B-tubule; IDA, inner dynein arm; a–e and g, inner dynein arm isoforms; I1α/ I1β/ ICLC, α- and β-heavy chain, and intermediate-light chain complex of I1 dynein; L1, L1 protrusion connecting between the linker and dynein g; ODA, outer dynein arm. Scale bar: 10 nm.

**Fig. S5.**
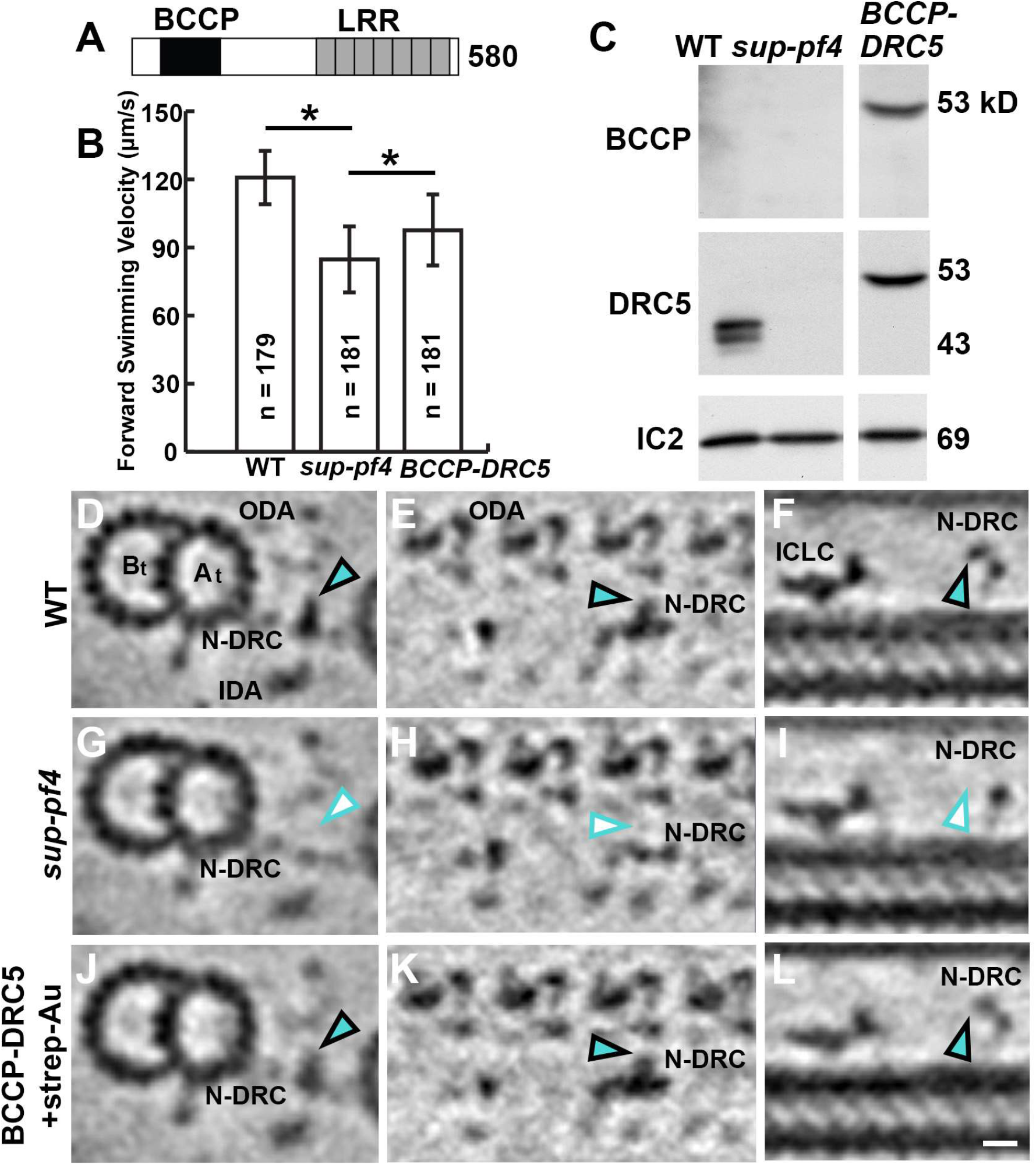
BCCP-DRC5 rescues the *sup-pf4* mutant phenotypes and restores the missing N-DRC structures, but the location of the N-terminal label of DRC5 could not be determined. **(A)** Diagram of the DRC5 polypeptide sequence with predicted domains indicated: LRR, leucine-rich repeat; BCCP, biotin carboxyl carrier protein (inserted near the N terminus of DRC5). **(B)** The forward swimming velocities of *sup-pf4* are shown relative to those of WT and the rescued *sup-pf4;N-BCCP-DRC5* strain. The *sup-pf4* was significantly slower (P < 0.05) than both WT and the *BCCP-DRC5* rescued strain. *BCCP-DRC5* was significantly faster than the mutant but slightly slower than WT (P < 0.05). **(C)** Western blot of axonemes isolated from WT, *sup-pf4*, and the *BCCP-DRC5* rescue strain was probed with antibodies against DRC subunits. Note the presence of a DRC5 band in the rescued strain that was assembled at WT levels but migrated at the size predicted for the BCCP-tagged protein. Antibodies against the IC2 subunit of the outer dynein arm served as loading control. **(D-L)** Tomographic slices of the averaged 96-nm axonemal repeats from wild type (D-F), *sup-pf4* (G-I) and the rescued *sup-pf4;N-BCCP-DRC5* strain (J-L) viewed in cross-sectional **(D, G, J)** and longitudinal view **(E, F, H, I, K, L).** Purified axonemes of BCCP-tagged DRC5 were labeled with strep-Au (1.4 nm nanogold). Electron densities corresponding to the L2 protrusion of the nexin-dynein regulatory complex (N-DRC) linker (cyan arrowheads in D-F) were missing in *sup-pf4* (white arrowheads in G-I), indicating the location of the DRC5/6 subcomplex, and were restored in the *BCCP-DRC5* rescued strain (cyan arrowheads in J-L). However, we did not observe any additional densities corresponding to the streptavidin-nanogold bound to the BCCP-DRC5 axonemes. Other labels: At, A-tubule; Bt, B-tubule; IDA, inner dynein arm; ODA, outer dynein arm; ICLC, intermediate and light chain complex. Scale bar: 10 nm.

**Table S1.**
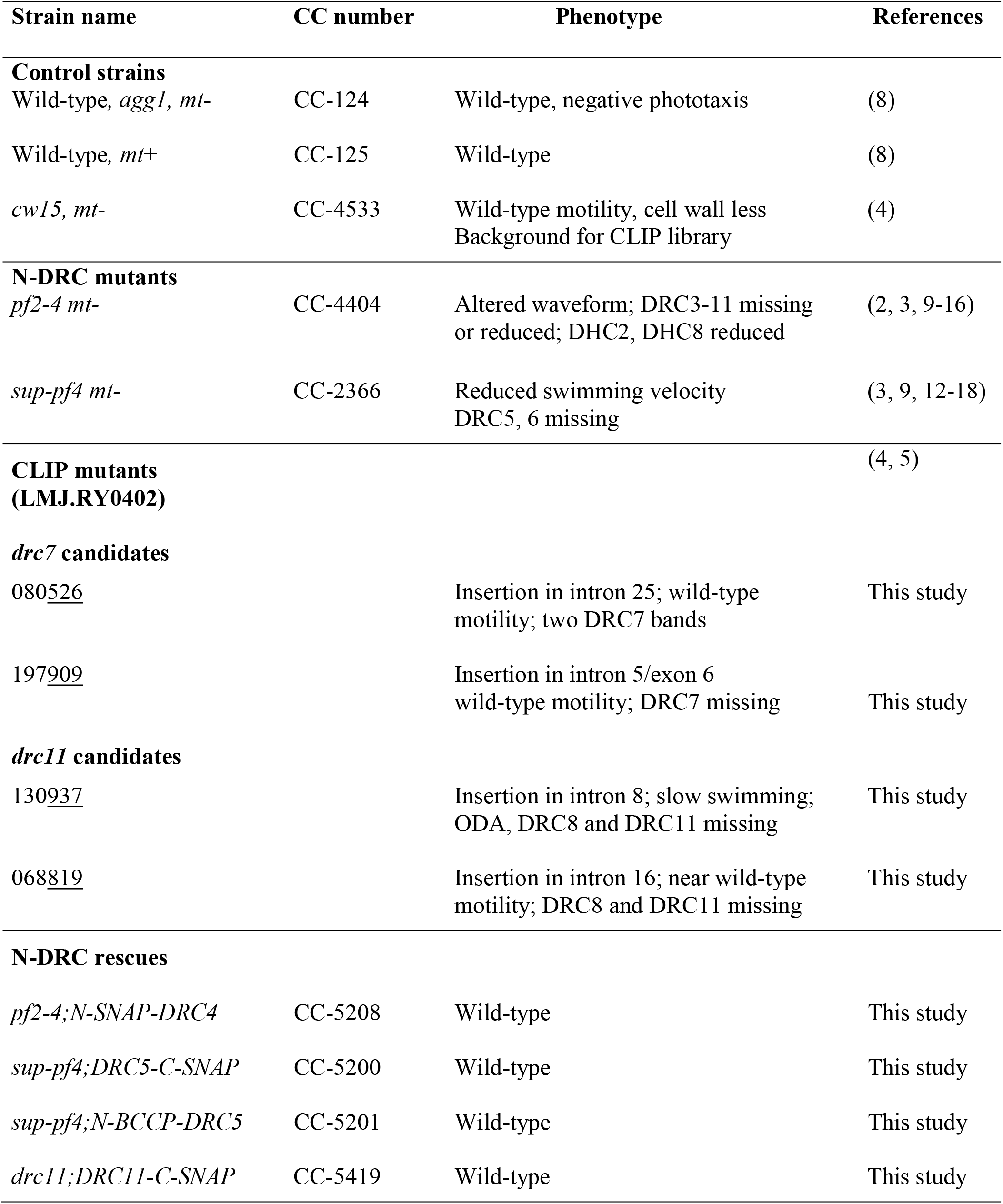
Strains used in this study.

**Table S2.** Image processing information for strains used in this study.

| Name | Strain | Tomograms included | Averaged repeats | Resolution<sup>a</sup> (nm) | Used in Figs. |
| --- | --- | --- | --- | --- | --- |
| WT<sup>b,d</sup> | *CC-125* | 19 | 2381 | 2.7 | 1, 3-5, 8, S2 |
| *drc7*<sup>b</sup> | *drc7-197909* | 11 | 1550 | 2.8 | 3 |
| *drc11*<sup>b</sup> | *drc11-068819* | 19 | 2544 | 2.6 | 4,5 |
| *drc11-937*<sup>b</sup> | *drc11-130937* | 9 | 1100 | 2.9 | S2 |
| WT<sup>c,d</sup> | *CC-125* | 25 | 3736 | 3.0 | 7, S3, S4, S5 |
| SNAP-DRC4<sup>c</sup> | *pf2-4;N-SNAP-DRC4* | 9 | 973 | 3.5 | 7, S3 |
| *sup-pf-4*<sup>c,d</sup> | *sup-pf-4* | 3 | 500 | 3.8 | S5 |
| DRC5-SNAP<sup>c</sup> | *sup-pf-4;DRC5-C-SNAP* | 3 | 364 | 4.0 | 7, S4 |
| BCCP-DRC5<sup>c</sup> | *sup-pf-4;N-BCCP-DRC5* | 8 | 766 | 3.6 | S5 |

<sup>a</sup> The 0.5 Fourier shell correlation criterion was used to estimate the resolution of the averaged repeats at the site of the doublet microtubule;
<sup>b</sup> Data recorded using Titan Krios TEM equipped with K2 and Volta-Phase-Plate;
<sup>c</sup> Data recorded using F30 with CCD;
<sup>d</sup> Includes previously published data: WT (19); *sup-pf-4* (15).

**Table S3.**
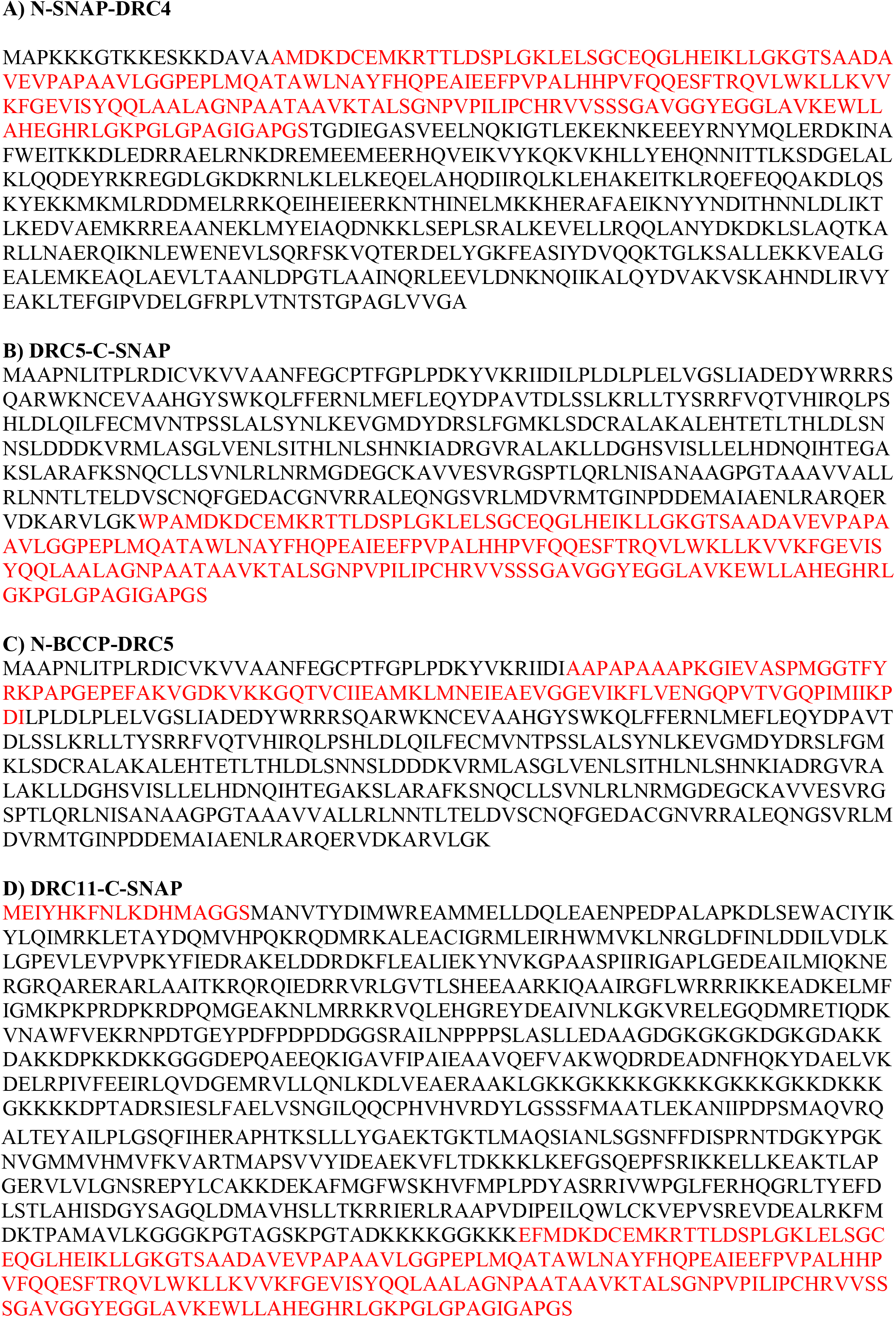
Predicted amino acid sequences of tagged DRC subunits. The amino acid residues of the DRC subunits are shown in black, and the novel amino acids introduced by the cloning vectors and epitope tags are shown in red.

**Table S4.**
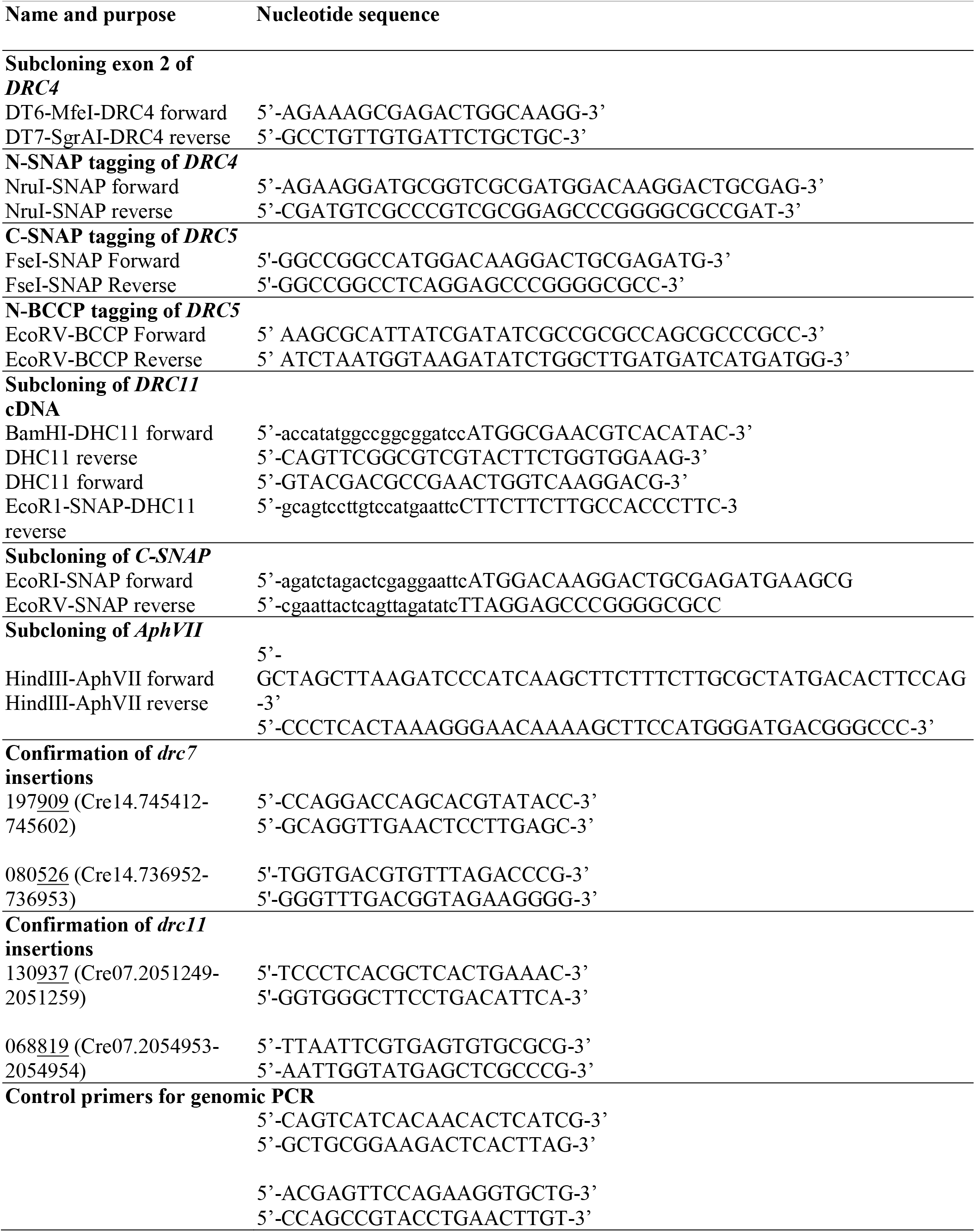
Oligonucleotide primers used in this study.

**Table S5.** Antibodies used in this study.

| Antibody | Host | Dilution (WB) | Reference or source |
| --- | --- | --- | --- |
| DRC1 fusion | Rabbit | 1:1000-10000 | (20) |
| DRC2 fusion | Rabbit | 1:1000-1:50000 | (3) |
| DRC3 peptide | Rabbit | 1:50-1:1000 | (3) |
| Gas8 fusion (DRC4) | Rabbit | 1:1000-50000 | (3) |
| DRC5 peptide | Rabbit | 1:1000 | (3) |
| DRC7 fusion | Rabbit | 1:1000-1:10000 | (3) |
| DRC8 peptide | Rabbit | 1:200-1:10000 | (3) |
| DRC11 fusion | Rabbit | 1:1000-50000 | (3) |
| IC2 (IC69) | Mouse | 1:10000-1:20000 | Sigma #D6168 |
| SNAP | Rabbit | 1:1000 | New England Biolabs #P9310S |
| RSP16 | Rabbit | 1:10000 | (21) |
| CCDC39/FAP59 | Rabbit | 1:10000 | Sigma #HPA035364 |
| AOX1 | Rabbit | 1:15000 | Agrisera #AS06-152 |

**Table S6.** iTRAQ protein ratios in WT (*cw15*) and *drc11-937* axonemes.

| **Protein** | **Peptides** | **WT/WT** | ***drc11*/WT** |
| --- | --- | --- | --- |
| **N-DRC** |  |  |  |
| DRC2 | 40 (58) | 1.01 | 0.99 |
| DRC3 | 57 (73) | 0.98 | 0.96 |
| DRC4 | 89 (106) | 1.02 | 0.98 |
| DRC5 | 44 (19) | 0.99 | 0.96 |
| DRC6 | 16 (26) | 0.97 | 0.96 |
| DRC7 | 140 (179) | 1.00 | 0.99 |
| DRC8 | 9 (10) | 1.01 | **0.07** |
| DRC9 | 49 (52) | 1.00 | 0.98 |
| DRC10 | 28 (41) | 1.03 | 0.98 |
| DRC11 | 60 (71) | 0.99 | **0.11** |
| **Outer Arm** |  |  |  |
| DHC13 | 442 (475) | 0.98 | **0.09** |
| DHC14 | 468 (535) | 0.97 | **0.09** |
| DHC15 | 306 (354) | 0.96 | **0.07** |
| DIC1 | 64 (83) | 1.01 | **0.10** |
| DIC2 | 60 (66) | 1.01 | **0.07** |
| DLU1/LC1 | 23 (24) | 1.00 | **0.11** |
| DLT2/LC2 | 19 (14) | 1.05 | **0.08** |
| DLX1/LC3 | 11 (13) | 0.99 | **0.07** |
| DLE1/LC4 | 9 (9) | 0.92 | **0.07** |
| DLX2/LC5 | 16 (20) | 0.99 | **0.09** |
| DLL2/LC6 | 15 (20) | 1.10 | **0.10** |
| DLR1/LC7a | 16 (13) | 0.97 | **0.38** |
| DLR2/LC7b | 22 (26) | 1.01 | **0.28** |
| DLL1/LC8 | 52 (53) | 1.08 | **0.69** |
| DLT1/LC9 | 15 (13) | 0.99 | **0.71** |
| DLL3/LC10 | 8 (7) | 0.97 | **0.07** |
| DCC1 | 59 (74) | 1.00 | **0.08** |
| DCC2 | 78 (96) | 1.00 | **0.10** |
| DCC3 | 17 (22) | 0.91 | **0.06** |
| **Others (<0.15)** |  |  |  |
| PRK1 | 10 (10) | 0.87 | **0.10** |
| NAGK1 | 7 (3) | 0.98 | **0.11** |
| Cre02.g095150 | 19 (14) | 0.97 | **0.11** |
| FMG-1B | 61 (46) | 0.99 | **0.14** |
| Cre03.g192201 | 5 (5) | 0.99 | **0.14** |

Samples of WT (*cw15*) and *drc11-937* axonemes were labeled with iTRAQ tags (technical replicates) and subjected to trypsin digestion and MS/MS. 1080 polypeptides were detected at a false discovery rate of 5%. Peptides indicate the number of peptides used for protein identification (95% confidence) followed by number used for iTRAQ quantification (in parenthesis). The amount of protein in each sample was compared to that present in WT to obtain a relative protein ratio. The WT/WT ratio indicates the variability in iTRAQ labeling and protein loading between technical replicates (typically less than 10%). The two mutant/WT ratios from the technical replicates were averaged. Shown here are the ratios for the N-DRC subunits, the ODA subunits, and other proteins whose ratios were less than 15% of WT. Those ratios that were statistically significant in both replicates are shown in bold.

**Movie S1.**
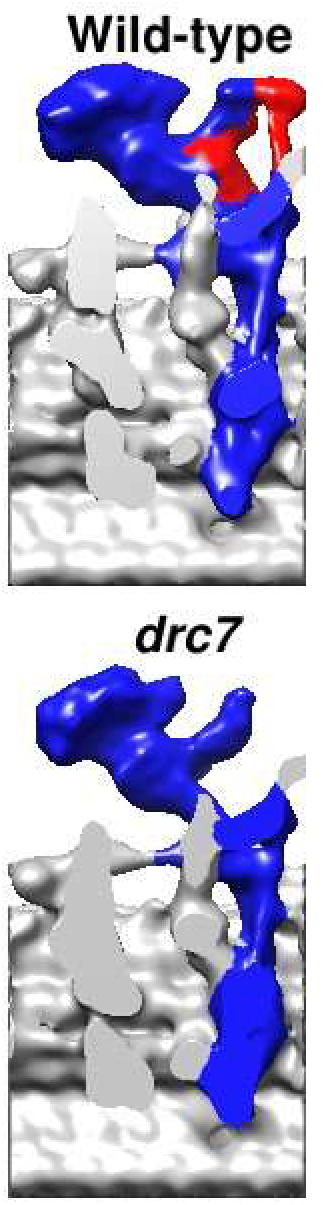
Three-dimensional visualization of the N-DRC from wild type and *drc7* in longitudinal bottom views. Animated 3D visualization compares the isosurface renderings of the nexin-dynein regulatory complex (N-DRC) structures from wild-type (top) and *drc7* (bottom) axonemes. The proximal end of the flagella is on the left. The red-colored densities represent the location of DRC7 at the OID linker, a portion of the distal lobe (dL), and the connection between the distal lobe and the L1 protrusion. Compare with Fig. 3.

**Movie S2.**
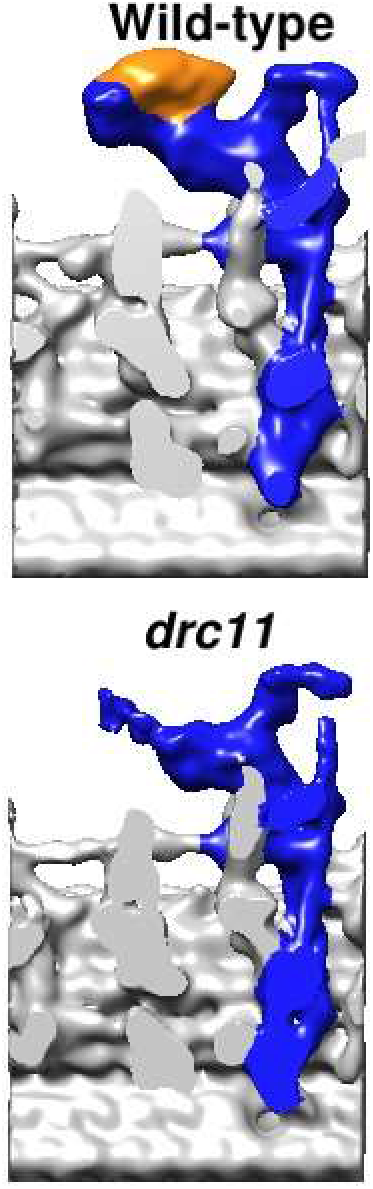
Three-dimensional visualization of the N-DRC from wild type and *drc11* in longitudinal bottom views. Animated 3D visualization compares the isosurface renderings of the nexin-dynein regulatory complex (N-DRC) structures from wild-type (top) and *drc11* (bottom) axonemes. The proximal end of the flagella is on the left. The orange-colored densities represent the location of the DRC8/11 sub-complex at the top portion of the proximal lobe. Compare with Fig. 4.

**Movie S3.**
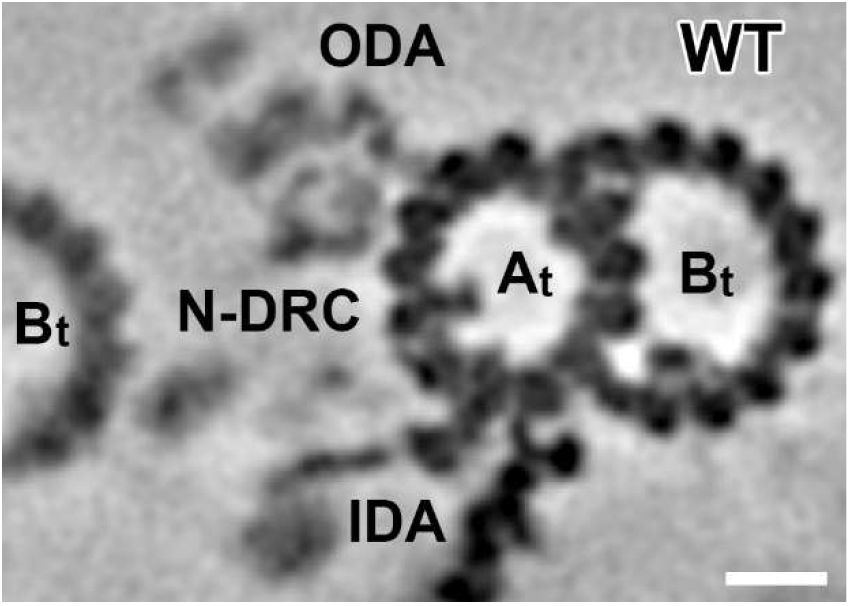
Tomographic slices of the averaged axonemal repeats from wild-type (100%) and *drc11* axonemes (100% and 3 class averages) show the proximal lobe of the nexin-dynein regulatory complex (N-DRC) in cross-sectional view. Note that in *drc11*, the proximal lobe density is reduced and blurry. A classification analysis of this density revealed three classes that vary in the positions of the proximal lobe between the rows of inner and outer dynein arms (IDA, ODA). Other labels: At, A-tubule; Bt, B-tubule. Compare with Fig. 5. Scale bar, 10 nm.

**Movie S4.**
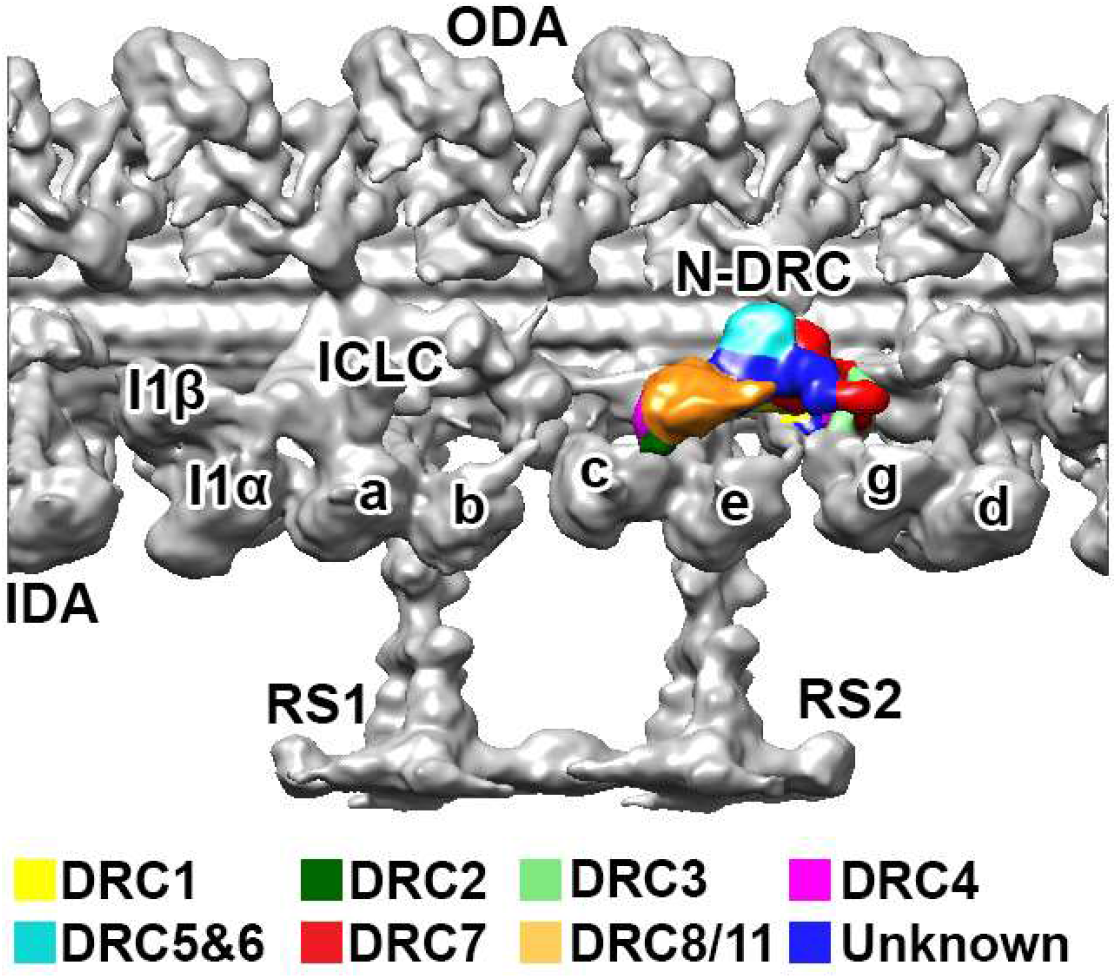
Three-dimensional localization of the DRC subunits and N-DRC interactions with neighboring structures. An animated 3D visualization shows an isosurface rendering of the averaged 96-nm axonemal repeat in wild type. At the beginning of the movie, proximal is on the left. The structural comparison between the wild-type average and deletion mutants of DRC subunits or mutant-rescues with tagged DRC subunits revealed the architecture of the nexin-dynein regulatory complex (N-DRC): the core scaffold extends along the entire length of the N-DRC and consists of DRC1 (yellow), DRC2 (dark green), and DRC4 (magenta); with this scaffold associate several functional subunits, i.e. DRC3 (light green), DRC5/6 subcomplex (cyan), DRC7 (red), and DRC8/11 subcomplex (orange). The composition of a few areas within the N-DRC remain unknown (dark blue). Other labels: At, A-tubule; Bt, B-tubule; IDA, inner dynein arm; ODA, outer dynein arm; RS, radial spoke; a–e and g, inner dynein arm isoforms; I1α/ I1β/ ICLC, α- and β-heavy chain, and intermediate-light chain complex of I1 dynein. Compare with Fig. 8.

